# Neurogenesis-mediated circuit remodeling reduces engram reinstatement and promotes forgetting

**DOI:** 10.1101/2023.10.10.561722

**Authors:** Axel Guskjolen, Jagroop Dhaliwal, Juan de la Parra, Jonathan R. Epp, Sangyoon Ko, Erin Chahley, Mitch de Snoo, Michela Solari, Prabdeep Panesar, Jason Snyder, Xin Duan, Joshua R. Sanes, Sheena A. Josselyn, Paul W. Frankland

## Abstract

Post-training increases in hippocampal neurogenesis are associated with forgetting of hippocampus-dependent memories in adult mice. This form of forgetting might be due to increased numbers of new neurons, remodeling of hippocampal circuitry or some combination of both. Here we tested the hypothesis that neurogenesis-mediated forgetting is caused by remodeling of hippocampal circuits by engineering mice in which adult-generated granule cells hypo- or hyper-integrate into hippocampal circuits. Using gene deletion, opto- and chemogenetic strategies, we find that hypo-integration of newborn neurons prevents post- training exercise-induced forgetting of contextual fear memories. Conversely, inducing hyper- integration of newborn neurons following contextual fear conditioning is sufficient to produce forgetting. Because these interventions did not affect survival of newborn neurons, these findings suggest that neurogenesis-mediated remodeling of hippocampal circuits represents a continuous and active form of interference that alters accessibility of engrams underlying hippocampal memories. Consistent with this, using engram-labeling approaches, we found that exercise-induced forgetting was associated with reduced engram reactivation.

## INTRODUCTION

The hippocampus is thought to automatically encode our experiences, allowing the present and very recent past to be continuously available for conscious recollection^1^. However, unless something remarkable happens, the vast majority of these every day experiences are ultimately forgotten^2^. Given the apparent pervasiveness of forgetting, there must exist active forgetting mechanisms that clear information from the hippocampus, as well as other neural systems^3–5^. Indeed, active forgetting processes are hypothesized to play an important role in optimizing mnemonic processing^6–8^. For example, forgetting may (i) remove outdated information and allow more flexible updating in changing environments, and (ii) prevent overfitting to specific past events and allow more accurate future predictions^6^.

Recent neurobiological studies in flies and rodents have begun to identify several distinct forms of active forgetting that are dissociable with respect to mechanism and timescale^4,6,9^. Our studies on the neurobiology of forgetting have focused on hippocampal neurogenesis^5^. In mice, environmental, pharmacological or genetic interventions that elevate levels of hippocampal neurogenesis immediately following learning induce forgetting of hippocampus-dependent memories^10–15^, but leave hippocampus-independent memories intact^10^. Conversely, interventions that decrease levels of hippocampal neurogenesis in the post-learning period reduce forgetting of hippocampus-dependent memories^10,11,15^. These forgetting effects are observed across a range of aversively- and appetitively-motivated, hippocampus-dependent paradigms^10–13^. Furthermore, post-training elevation of hippocampal neurogenesis induces forgetting in other rodent species, including rats, degus and guinea pigs^10,16^.

While these experiments establish that post-training changes in hippocampal neurogenesis levels bidirectionally modulate the stability of encoded hippocampal memories, how this occurs remains unclear. As new neurons integrate into established hippocampal circuits, they gradually form input and output connections, starting around 16 days^17^. This is a competitive process, whereby new synaptic connections may co-exist with, or replace, existing synaptic connections^18–21^. Since successful recall of hippocampal memories depends on reinstatement of neural ensembles that were active during initial learning (i.e., engram reactivation), then changes in circuit architecture as a consequence of hippocampal neurogenesis might reduce the probability of engram reactivation and lead to forgetting^5,22^. Indeed, computational models examining how neurogenesis impacts the stability of stored information indicate that forgetting stems from a change in network connection patterns. While adding new neurons produces forgetting in these network models, changing connectivity patterns of new neurons, without changing their number, is also sufficient to induce memory loss^23–25^.

Here we test whether changes in connectivity, rather than overall increases in new neuron number, are responsible for forgetting and whether this form of forgetting reduces engram reinstatement. To do this we developed transgenic, chemo- and optogenetic strategies to modulate how new neurons integrate into adult, hippocampal circuits (i.e., without affecting their survival or overall levels of neurogenesis). In the first approach, we engineered mice in which new neurons *hypo*-integrate into hippocampal circuits during adulthood. By attenuating integration, and associated changes in network connectivity, we predicted that forgetting produced by post-training increases in hippocampal neurogenesis would be reduced or even prevented. In the second approach, we engineered mice in which new neurons *hyper*-integrate into hippocampal circuits during adulthood. In these mice, we predicted hyper-integration of new neurons, and associated changes in network connectivity, would be sufficient to induce forgetting in the absence of any changes in overall levels of adult neurogenesis. Finally, we assessed whether neurogenesis-mediated forgetting is associated with reduced engram reinstatement, and whether artificial chemogenetic engram reactivation can help restore otherwise forgotten memories.

## RESULTS

### Elimination of Rac1 attenuates dendritic growth, spine density and large mossy fiber terminal (LMT) density in newborn granule cells

Post-training exercise increases hippocampal neurogenesis and induces forgetting of hippocampus-dependent memories^10–12^. We first asked whether restricting the extent to which adult-generated neurons integrate into hippocampal circuits might reduce or even eliminate forgetting induced by post-training exercise. A previous study indicated that the small Rho GTPase, Rac1, regulates the integration of adult-generated granule cells into hippocampal circuits^26^. In this study, neural precursor cell (NPC) specific Rac1 deletion or dominant-negative Rac1 over-expression reduced dendritic growth and spine density of adult-generated granule cells, measured one month later. Importantly, in either case, Rac1-deficiency did not lead to a reduction in the number of adult-generated granule cells surviving to one month, suggesting that Rac1 is not required for the initial stages of neuronal development of adult-generated granule cells.

In order to confirm that elimination of Rac1 attenuates integration of adult-generated granule cells without affecting survival, we combined viral and genetic loss-of-function strategies in adult mice. We injected retrovirus (RV) expressing GFP and Cre-recombinase (i.e., RV-Cre^+^) or GFP alone (i.e., RV-Cre^-^) into the dentate gyrus (DG) of adult floxed Rac1 (Rac1^f/f^) mice (Fig. 1a).

**Figure 1.**
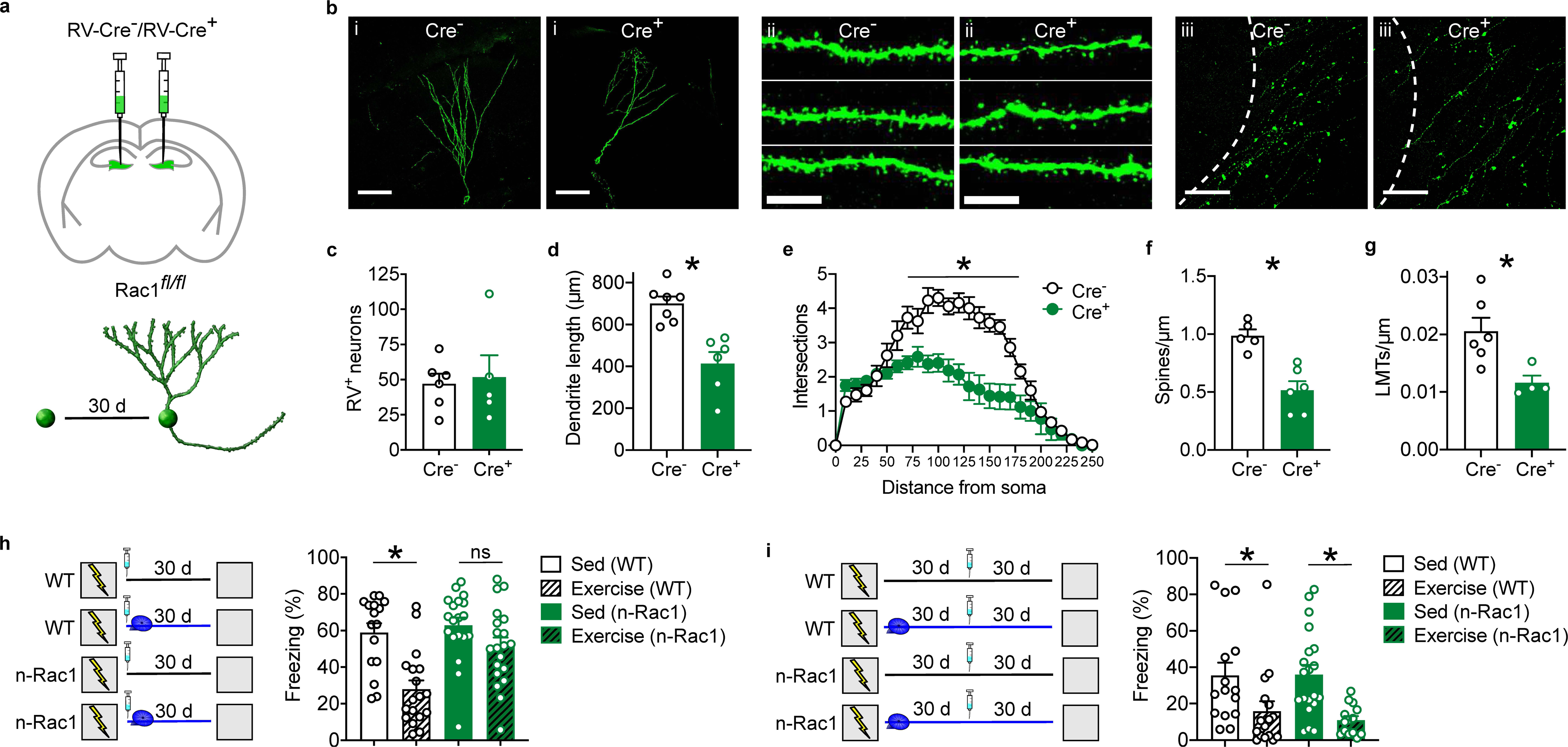
Rac1 deletion in NPCs induces hypo-integration and prevents exercise-induced forgetting. (**a**) Experimental design. Rac1^f/f^ mice were micro-infused with GFP-tagged RV-Cre^-^ or RV-Cre^+^, and the morphology of infected adult-generated granule cells was assessed 30 d later. (**b**) Representative images from hippocampus showing (i) dendritic morphology (scale bar = 50 µm), (ii) dendritic segments (scale bar = 10 µm) and (iii) LMTs (scale bar = 50 µm) of Cre^-^ and Cre^+^ infected adult-generated granule cells. Rac1 deletion (Cre^+^ cf. Cre^-^) did not affect (**c**) survival of adult-generated granule cells (*t*_9_ = 0.30, *P* = 0.77), but induced hypo-integration, as reflected in reduced (**d**) total dendrite length (*t*_11_ = 4.59, *P* < 0.001), (**e**) dendritic complexity (ANOVA, main group effect: *F*_1,11_ = 17.87, *P* < 0.005; main distance effect: *F*_24,264_ = 50.77, *P* < 0.001; group × distance interaction: *F*_24,264_ = 9.12, *P* < 0.001), (**f**) spine density (*t*_9_ = 4.75, *P* < 0.005) and (**g**) LMT density (*t*_8_ = 2.89, *P* < 0.01). (**h**) (left) Experimental design. WT and n-Rac1 mice were trained in contextual fear conditioning. During the 30 d retention delay mice either had access to an exercise wheel (‘Exercise’ groups) or were housed conventionally (‘Sed’ [sedentary] groups). One day after training mice were treated with TAM to induce Rac1 deletion in n-Rac1 mice. (right) Post-training exercise induced forgetting in WT but not n-Rac1 mice (ANOVA, main genotype effect: *F*_1,69_ = 8.44, *P* < 0.005; main exercise effect: *F*_1,69_ = 20.15, *P* < 0.001; genotype × exercise interaction: *F*_1,69_ = 4.43, *P* < 0.05). (**i**) Experimental design. WT and n-Rac1 mice were trained in contextual fear conditioning, and tested 60 d later. Mice either had access to an exercise wheel (‘Exercise’ groups) or were housed conventionally (‘Sed’ groups) following training. Mice were treated with TAM 30 d (rather than one day) following training to induce Rac1 deletion in n-Rac1 mice. (right) Post-training exercise induced forgetting in both WT and n-Rac1 mice (ANOVA, main exercise effect: *F*_1,61_ = 16.70, *P* < 0.001; no genotype effect: *F*_1,61_ = 0.16, *P* = 0.68; no genotype × exercise interaction: *F*_1,61_ = 0.24, *P* = 0.62).

Thirty days later, numerous infected granule cells were identified in the innermost layers of the granule cell layer in the DG (Fig. 1b). Similar numbers of control (Cre^-^) and Rac1-deficient (Cre^+^) infected granule cells were identified (Fig. 1c), suggesting that Rac1 elimination did not affect survival of adult-generated granule cells (for convergent evidence using BrdU and DCX labeling, see Extended Data Fig. 1a-d). However, Rac1-deficient neurons had reduced total dendritic length (Fig. 1d), and less complex dendritic arbors (Fig. 1e). Moreover, both spine density (Fig. 1f) and density of LMTs in CA3 (Fig. 1g) were reduced, suggesting both input and output connectivity of newborn neurons is attenuated following Rac1 elimination.

### Reduced integration of Rac1-deficient granule cells prevents forgetting

If neurogenesis-dependent forgetting depends on circuit remodeling, then restricting the ability of new neurons to form input and output connections should prevent forgetting. Our analyses confirmed that elimination of Rac1 results in hypo-integration of adult-generated granule cells, without affecting neuronal survival^26^, and therefore manipulating Rac1 in adult-generated neurons provide an ideal approach to test this hypothesis.

We crossed Rac1^f/f^ mice with mice expressing a tamoxifen (TAM) inducible Cre-recombinase under the nestin promotor that is active in NPCs (nestin-Cre^ERT2^ mice). In adult offspring from this cross that express both transgenes (n-Rac1 mice), TAM treatment induces deletion of Rac1 from nestin^+^ cells. Adult n-Rac1 and their littermate controls were placed in a novel training context and received 3 footshocks. They were treated with TAM one day later, and then mice were either allowed continuous access to an exercise wheel in their home cage (‘exercise’ groups) or housed conventionally (‘sedentary’ groups). Contextual fear memory was assessed 30 days following training by replacing mice in the training context and measuring freezing behavior^27^ (Fig. 1h).

In this test, control mice that exercised following training showed reduced levels of conditioned fear compared to sedentary controls. In contrast, this exercise-induced forgetting was eliminated in n-Rac1 mice, with both sedentary and exercise n-Rac1 mice freezing at similar levels to the WT sedentary group (Fig. 1h). Therefore, restricting integration of adult-generated granule cells prevented forgetting that is normally produced by post-training elevation of hippocampal neurogenesis.

By manipulating how adult-generated neurons integrate into hippocampal circuits, these data support the hypothesis that forgetting is due to circuit remodeling. However, altered hippocampal neurogenesis may be associated with increased anxiety-like behaviors in mice^28^. Therefore, another possibility is that perturbations in neurogenesis increase anxiety-like states in the n-Rac1 mice, which, in turn, promote freezing behavior in the memory test. We addressed this potential confound in two additional experiments.

First, we delayed Rac1 deletion until after forgetting had occurred. In this experiment, WT and n-Rac1 mice were trained, and given home cage access to an exercise wheel (‘exercise’ groups) or housed conventionally (‘sedentary’ groups) for 30 days. The mice were then treated with TAM, and contextual fear was assessed 30 days later (i.e., ∼8.5 weeks following initial training) (Fig. 1i). Since 4 weeks of exercise is sufficient to induce forgetting^12^, we reasoned that any changes in freezing levels would be attributed to non-specific effects of Rac1 deletion. In the memory test, exercise mice exhibited lower levels of freezing compared to sedentary mice, as expected. Importantly, Rac1 deletion did not alter freezing levels, suggesting that elimination of Rac1 from NPCs does not non-specifically alter anxiety-like states. Moreover, these results suggest that, beyond a critical window, exercise-induced forgetting effects are not reversible by manipulating hippocampal neurogenesis.

In a second experiment, we assessed anxiety-related behaviors by testing separate cohorts of n-Rac1 and littermate control mice in the open field (Extended Data Fig. 1e-f). In the open field, total distance travelled was equivalent in n-Rac1 and control mice. Moreover, while mice spent more time in the periphery of the open field, this spatial bias did not differ between groups, suggesting that Rac1 deletion did not alter anxiety-related behaviors.

### Deletion of Rac1 from mature excitatory cells does not block forgetting

Experience-dependent remodeling of mature, excitatory cells may also occur^29–31^. In order to evaluate whether these effects were specific to Rac1 deletion from immature cells, we examined the impact of Rac1 deletion on the morphology of mature excitatory DG neurons by crossing Rac1^f/f^ mice with mice expressing a tamoxifen (TAM) inducible Cre-recombinase under the α-CaMKII promotor (α-CaMKII-Cre^ERT2^ mice). Adult offspring from this cross expressing both transgenes (c-Rac1 mice) and their littermate controls were treated with TAM. HSV-GFP (which infects mature, excitatory neurons) was infused into the DG 26 days later. On day 30, the morphology of DG granule cells was examined, as previously (Fig. 2a). In contrast to newly- generated neurons, Rac1 deletion had no impact on morphology (Fig. 2b). Dendritic length, complexity, spine and LMT density were similar in c-Rac1 and control mice (Fig. 2c-f). This indicates that Rac1 deletion produces distinct morphological phenotypes in immature vs. mature granule cells.

**Figure 2.**
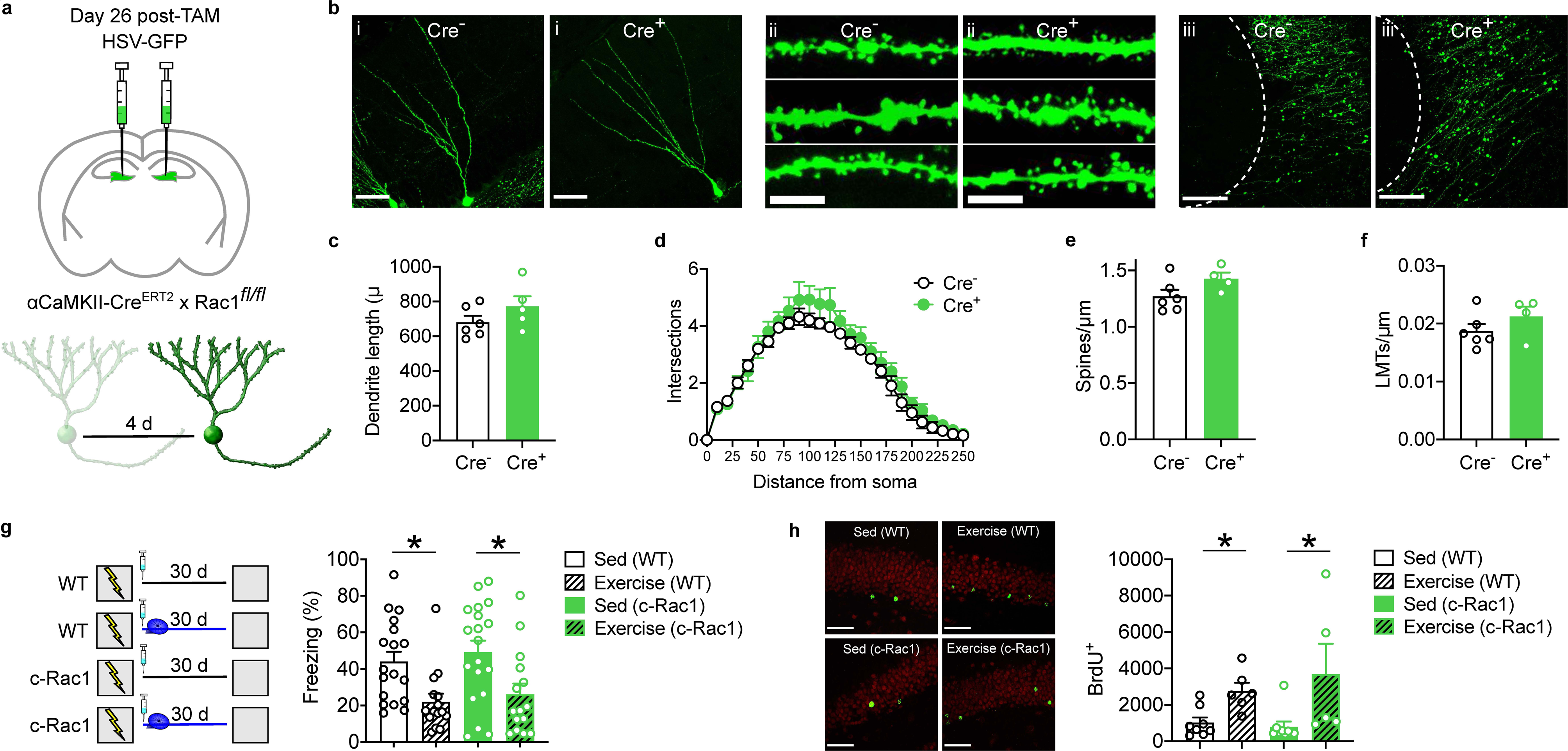
Rac1 deletion in mature neurons does not impact morphology or prevent exercise-induced forgetting. (**a**) Experimental design. c-Rac1 mice were treated with TAM, and then micro-infused with HSV expressing GFP 26 d later. The morphology of infected cells was assessed 4 d later. (**b**) Representative images from hippocampus showing (i) dendritic morphology (scale bar = 50 µm), (ii) dendritic segments (scale bar = 10 µm) and (iii) LMTs (scale bar = 50 µm) of infected cells. Rac1 deletion (Cre^+^ cf. Cre) did not affect (**c**) total dendrite length (*t*_9_ = 1.40, *P* = 0.20), (**d**) dendritic complexity (ANOVA, main group effect: *F*_1,9_ = 1.73, *P* = 0.22; main distance effect: *F*_27,243_ = 114.40, *P* < 0.0001; group × distance interaction: *F*_27,243_ = 0.75, *P* = 0.80), (**e**) spine density (*t*_8_ = 1.85, *P* = 0.10) nor (**f**) LMT density (*t*_8_ = 1.23, *P* = 0.25). (**g**) (left) Experimental design. WT and c-Rac1 mice were trained in contextual fear conditioning. During the 30 d retention delay, mice either had access to an exercise wheel (‘Exercise’ groups) or were housed conventionally (‘Sed’ groups). One day after training mice were treated with TAM to induce Rac1 deletion in mature neurons. (right) Post-training exercise induced forgetting in both WT and c-Rac1 mice (ANOVA, main exercise effect: *F*_1,63_ = 16.14, *P* < 0.001; no genotype effect: *F*_1,63_ = 0.67, *P* = 0.42; no genotype × exercise interaction: *F*_1,63_ = 0.01, *P* = 0.92). (**h**) (left) Representative images showing BrdU-labeled cells in DG (scale bar = 50 µm). (right) Exercise increased BrdU^+^ cells in both WT and c-Rac1 mice (ANOVA, main exercise effect: *F*_1,24_ = 11.90, *P* < 0.005; no genotype effect: *F*_1,24_ = 0.26, *P* = 0.61; no genotype × exercise interaction: *F*_1,24_ = 0.74, *P* = 0.40).

We next examined whether Rac1 deletion from mature excitatory neurons throughout the brain would prevent exercise-induced forgetting. c-Rac1 mice and their littermate controls were trained in contextual fear conditioning. They were treated with TAM one day later, and then allowed continuous access to an exercise wheel in their home cage (‘exercise’ groups) or housed conventionally (‘sedentary’ groups). Exercise reduced freezing levels in both control and c-Rac1 mice in the test 30 days following training, indicating that deletion of Rac1 throughout the brain does not affect neurogenesis-mediated forgetting (Fig. 2g). Moreover, Rac1 deletion from mature, excitatory cells did not impact levels of hippocampal neurogenesis, as assessed by BrdU labeling (Fig. 2h).

### Elimination of Cdh9 attenuates dendritic growth, spine density and LMT density in newborn granule cells

Cadherin-9 (Cdh9) is expressed in DG granule cells and CA3 pyramidal neurons and plays a key role in the formation of mossy fiber-CA3 synapses in the developing hippocampus. In particular, knockdown of Cdh9 from DG granule cells impairs formation of LMTs, leading to reductions in both number and size of LMTs, without affecting axon growth^32^. We reasoned that elimination of Cdh9 from NPCs in adult mice would similarly impact integration of adult- generated granule cells, with most pronounced impact on output connectivity (e.g., LMT density and size).

In floxed Cdh9 (Cdh9^f/f^) we injected GFP-tagged RV-Cre^+^ or RV-Cre^-^ into the DG. Thirty days later, we compared morphology of control (Cre^-^) vs. Cdh9-deficient (Cre^+^) granule cells (Fig. 3a). Infected granule cells were identified in the innermost layers of the granule cell layer in the DG (Fig. 3b), with no difference in number suggesting that Cdh9 deletion does not affect survival (Fig. 3c) (for convergent evidence using BrdU and DCX labeling, see Extended Data Fig. 2a-d). While Cdh9-deficiency did not affect dendritic complexity or spine density (Fig.3d-f), both LMT density and size were reduced in Cdh9-deficient cells (Fig. 3g-h). Therefore, similar to the developing hippocampus^32^, the loss of Cdh9 preferentially impacts the development of DG-CA3 connectivity.

**Figure 3.**
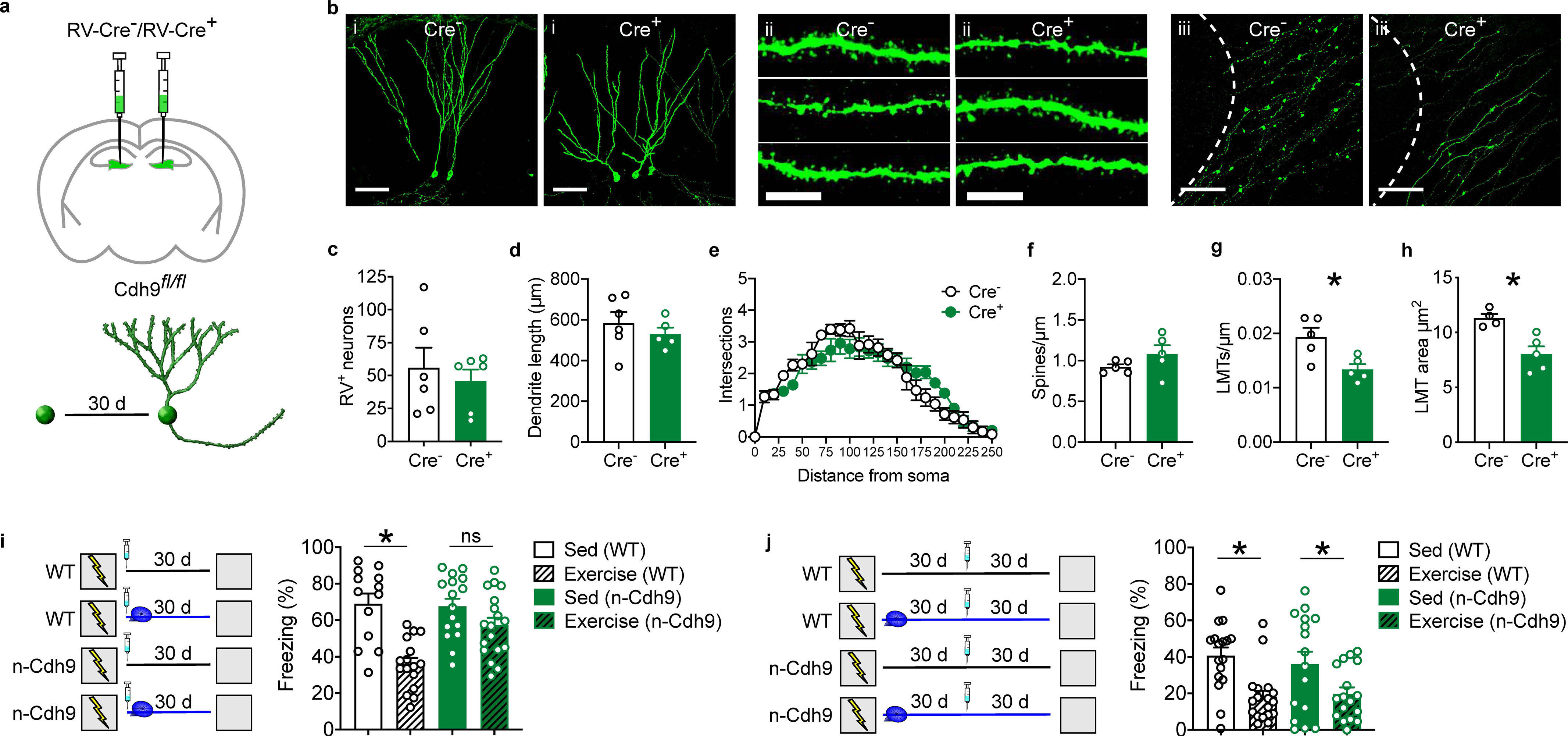
Cdh9 deletion in NPCs induces hypo-integration and prevents exercise-induced forgetting. (**a**) Experimental design. Cdh9^f/f^ mice were micro-infused with GFP-tagged RV-Cre^-^ or RV-Cre^+^, and the morphology of infected adult-generated granule cell was assessed 30 d later. (**b**) Representative images from hippocampus showing (i) dendritic morphology (scale bar = 50 µm), (ii) dendritic segments (scale bar = 10 µm) and (iii) LMTs (scale bar = 50 µm) of infected adult-generated granule cells. Cdh9 deletion (Cre^+^ cf. Cre^-^) did not affect (**c**) survival of adult-generated granule cells (*t*_10_ = 0.57, *P* = 0.58), (**d**) total dendrite length (*t*_9_ = 0.81, *P* = 0.44), (**e**) dendritic complexity (ANOVA, no main group effect: *F*_1,9_ = 0.03, *P* = 0.87; main distance effect: *F*_24,216_ = 10.27, *P* < 0.001; group × distance interaction: *F*_24,216_ = 2.77, *P* < 0.001), (**f**) nor spine density (*t*_8_ = 1.43, *P* = 0.19). Cdh9 deficiency reduced LMT (**g**) density (*t*_8_ = 3.03, *P* < 0.05) and (**h**) size (*t*_7_ = 3.77, *P* < 0.01). (**i**) (left) Experimental design. WT and n-Cdh9 mice were trained in contextual fear conditioning. During the 30 d retention delay mice either had access to an exercise wheel (‘Exercise’ groups) or were housed conventionally (‘Sed’ groups). One day after training mice were treated with TAM to induce Cdh9 deletion in n-Cdh9 mice. (right) Post- training exercise induced forgetting in WT but not n-Cdh9 mice (ANOVA, main genotype effect: *F*_1,59_ = 5.23, *P* < 0.05; main exercise effect: *F*_1,59_ = 24.94, *P* < 0.001, genotype × exercise interaction: *F*_1,59_ = 6.85, *P* < 0.05). (**j**) Experimental design. (left) WT and n-Cdh9 mice were trained in contextual fear conditioning, and tested 60 d later. Mice either had access to an exercise wheel (‘Exercise’ groups) or were housed conventionally (‘Sed’ groups) following training. Mice were treated with TAM 30 d (rather than one day) following training to induce Cdh9 deletion in n-Cdh9 mice. (right) Post-training exercise induced forgetting in both WT and n-Cdh9 mice (ANOVA, main exercise effect: *F*_1,47_ = 13.17, *P* < 0.001; no genotype effect: *F*_1,47_ = 0.01, *P* = 0.91; no genotype × exercise interaction: *F*_1,47_ = 0.09, *P* = 0.77).

### Reduced integration of Cdh9-deficient granule cells prevents forgetting

Since loss of Cdh9 promotes hypo-integration of adult-generated granule cells, without globally altering levels of adult neurogenesis, n-Cdh9 mice (nestin-Cre^ERT2^ × Cdh9^f/f^) provide a second approach to test our hypothesis that restricting integration of adult-generated granule cells into hippocampal circuits will prevent forgetting induced by post-training exercise. Accordingly, n- Cdh9 and their littermate controls were trained in contextual fear conditioning and tested 30 days later. During the retention delay, mice either had access to an exercise wheel in their home cage or were housed conventionally. As expected, post-training exercise induced forgetting in control mice. However, exercise-induced forgetting was not observed in the n-cdh9 mice (Fig. 3i).

The effects of deleting Cdh9 from neural progenitor cells on forgetting were temporally specific. Delaying Cdh9 deletion until after exercise-induced forgetting had occurred did not affect subsequent levels of conditioned fear (Fig. 3j). These data indicate that neurogenesis- dependent remodeling in the first 4 weeks following training is responsible for the forgetting effects. Moreover, they indicate that altered circuit integration of adult-generated granule cells in n-Cdh9 mice does not non-specifically promote freezing behavior by, for example, increasing anxiety. Consistent with this, n-Cdh9 mice did not display increased anxiety-like behavior in the open field task (Extened Data Fig. 2e-f). Finally, anatomical and behavioral analyses in c-Cdh9 (α-CaMKII-Cre^ERT2^ × Cdh9^f/f^) mice indicated that these effects are restricted to newborn neurons. Deletion of Cdh9 from mature excitatory neurons did not affect neuronal morphology (Fig. 4a-f), prevent forgetting associated with post-training exercise (Fig. 4g) or alter adult neurogenesis (Fig. 4h)

**Figure 4.**
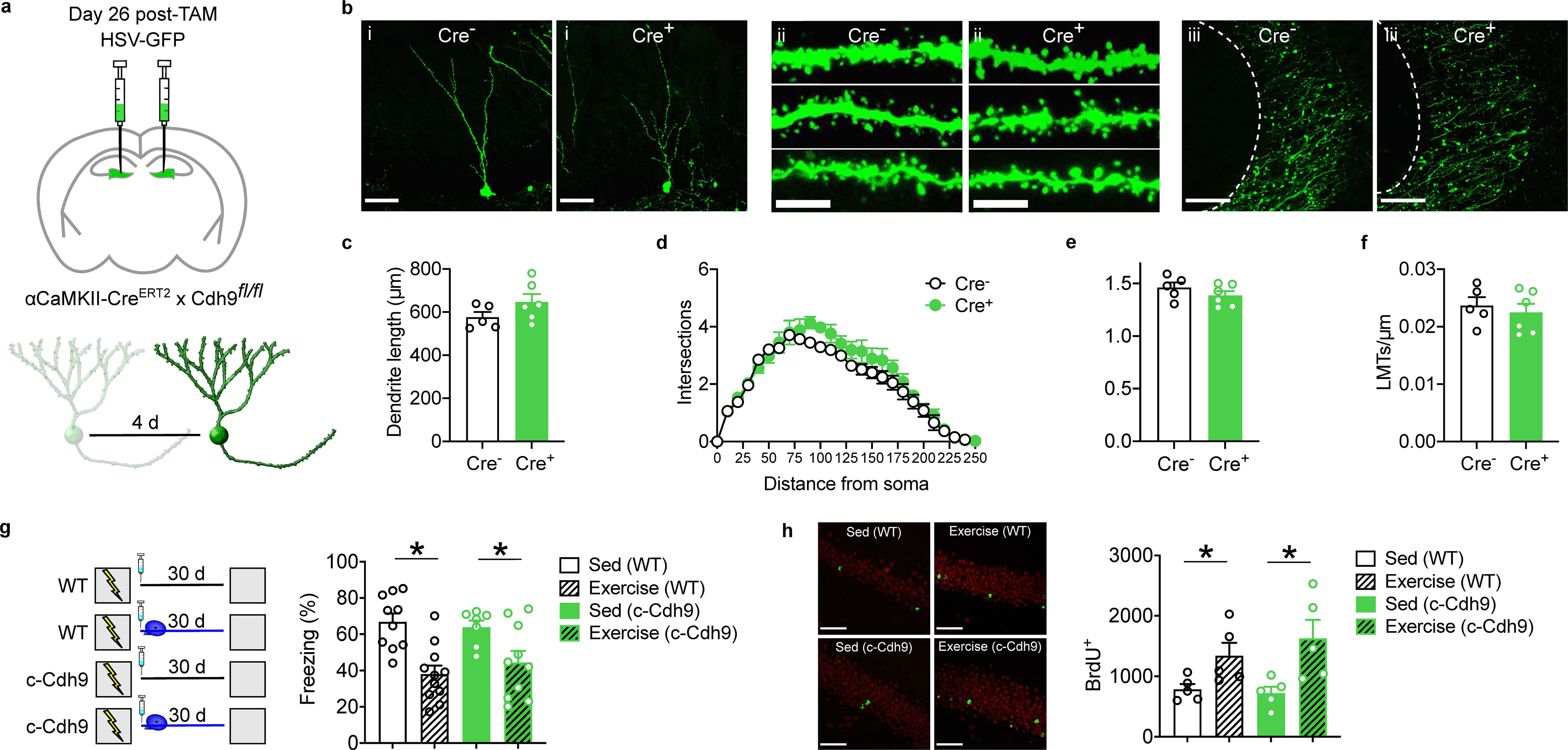
Cdh9 deletion in mature neurons does not impact morphology or prevent exercise-induced forgetting. (**a**) Experimental design. c-Cdh9 mice were treated with TAM, and then micro-infused with HSV expressing GFP 26 d later. The morphology of infected cells was assessed 4 d later. (**b**) Representative images from hippocampus showing (i) dendritic morphology (scale bar = 50 µm), (ii) dendritic segments (scale bar = 10 µm) and (iii) LMTs (scale bar = 50 µm) of infected cells. Cdh9 deletion (Cre^+^ cf. Cre) did not affect (**c**) total dendrite length (*t*_9_ = 1.54, *P* = 0.16), (**d**) dendritic complexity (ANOVA, no main group effect: *F*_1,9_ = 2.51, *P* = 0.15; main distance effect: *F*_24,216_ = 79.07, *P* < 0.0001; no group × distance interaction: *F*_24,216_ = 1.02, *P* = 0.44), (**e**) spine density (*t*_9_ = 1.17, *P* = 0.27) nor (**f**) LMT density (*t*_9_ = 0.56, *P* = 0.59). (**g**) (left) Experimental design. WT and c-Cdh9 mice were trained in contextual fear conditioning. During the 30 d retention delay, mice either had access to an exercise wheel (‘Exercise’ groups) or were housed conventionally (‘Sed’ groups). One day after training mice were treated with TAM to induce Cdh9 deletion in mature neurons. (right) Post-training exercise induced forgetting in both WT and c-Cdh9 mice (ANOVA, main exercise effect: *F*_1,34_ = 21.08, *P* < 0.001; no genotype effect: *F*_1,34_ = 0.11, *P* = 0.74; no genotype × exercise interaction: *F*_1,34_ = 0.80, *P* = 0.38). (**h**) (left) Representative images showing BrdU-labeled cells in DG (scale bar = 50 µm). (right) Exercise increased BrdU^+^ cells in both WT and c-Cdh9 mice (ANOVA, main exercise effect: *F*_1,16_ = 13.37, *P* < 0.005; no genotype effect: *F*_1,16_ = 0.31, *P* = 0.58; no genotype × exercise interaction: *F*_1,16_ = 0.76, *P* = 0.40).

### Elimination of semaphorin 5A promotes dendritic growth, increases spine density and LMT density in newborn granule cells

We next asked whether promoting, rather than restricting, integration of adult-generated neurons would be sufficient to induce forgetting. Our approach built on a previous study that showed that deletion of semaphorin 5A (Sema5A) increases spine density on adult-generated granule cells in the hippocampus, without affecting their survival^33^. In floxed Sema5A mice (Sema5A^f/f^) we injected GFP-tagged RV-Cre^+^ or RV-Cre^-^ into the DG and, 30 days later, compared morphology of control (Cre^-^) vs. Sema5A-deficient (Cre^+^) granule cells (Fig. 5a-b). Similar numbers of infected granule cells were identified in the innermost layers of the granule cell layer in the DG in both groups (Fig. 5c), indicating that Sema5A deletion does not affect survival (for convergent evidence using BrdU and DCX labeling, see Extended Data Fig. 3a-d). However, Sema5A deletion increased total dendrite length and dendritic complexity (especially at higher order branches) (Fig. 5d-e). Spine and LMT density were also elevated in Sema5A- deficient cells (Fig. 5f-g), indicating that Sema5A-deficiency impacts the development of input and output connectivity of adult-generated granule cells.

**Figure 5.**
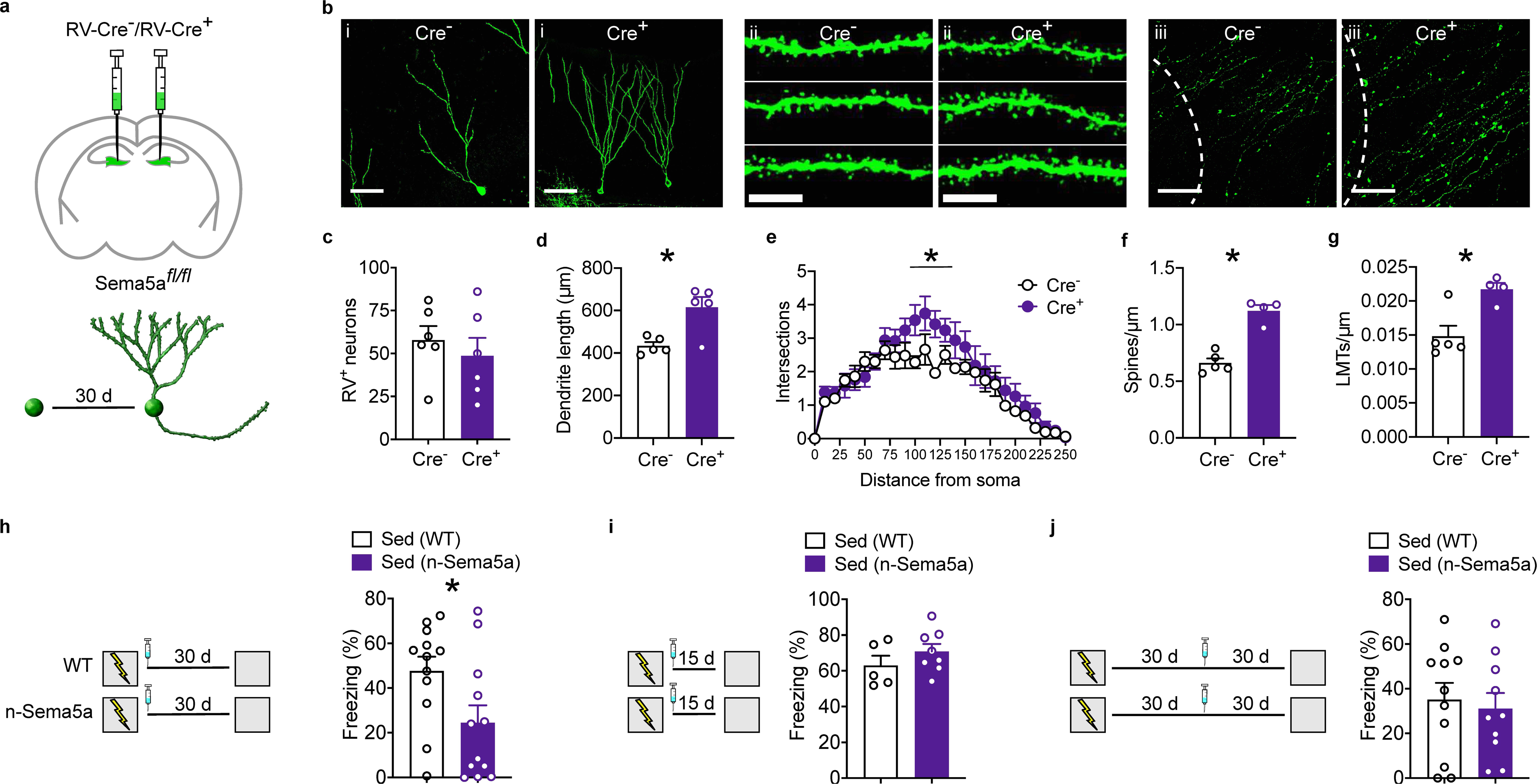
Sema5A deletion in NPCs induces hyper-integration and forgetting. (**a**) Experimental design. Sema5A^f/f^ mice were micro-infused with GFP-tagged RV-Cre^-^ or RV-Cre^+^, and the morphology of infected adult-generated granule cell was assessed 30 days later. (**b**) Representative images from hippocampus showing (i) dendritic morphology (scale bar = 50 µm), (ii) dendritic segments (scale bar = 10 µm) and (iii) LMTs (scale bar = 50 µm) of infected adult-generated granule cells. Sema5A deletion (Cre^+^ cf. Cre^-^) did not affect (**c**) survival of adult- generated granule cells (*t*_10_ = 0.69, *P* = 0.51), but increased (**d**) total dendrite length (*t*_9_ = 3.51, *P* < 0.01), (**e**) dendritic complexity (ANOVA, main group effect: *F*_1,8_ = 3.08, *P* = 0.12; main distance effect: *F*_24,192_ = 4.63, *P* < 0.0001; group × distance interaction: *F*_24,192_ = 1.39, *P* = 0.12), (**f**) spine density (*t*_7_ = 7.29, *P* < 0.0005), and (**g**) LMT density (*t*_7_ = 3.54, *P* < 0.005). (**h**) (left) Experimental design. WT and n-Sema5A mice were trained in contextual fear conditioning and tested 30 d later. One day after training mice were treated with TAM to induce Sema5A deletion in n- Sema5A mice. (right) Sema5A deletion induced forgetting (*t*_22_ = 2.31, *P* < 0.05). (**i**) (left) Experimental design. WT and n-Sema5A mice were trained in contextual fear conditioning and tested 15 d later. One day after training mice were treated with TAM to induce Sema5A deletion. (right) WT and i-Sema5A mice exhibited similar levels of freezing (*t*_11_ = 1.16, *P* = 0.27). (**j**) (left) Experimental design. WT and n-Sema5A mice were trained in contextual fear conditioning, and tested 60 d later. Mice were treated with TAM 30 d (rather than one day) following training to induce Sema5A deletion. (right) WT and n-Sema5A mice exhibited similar levels of freezing (*t*_16_ = 0.09, *P* = 0.38).

### Increased integration of Sema5A-deficient granule cells induces forgetting

We next tested whether induction of this hyper-integration phenotype would be sufficient to induce forgetting of contextual fear memories. To do this we crossed Sema5A^f/f^ mice with nestin-Cre^ERT2^ mice. Adult offspring from this cross expressing both transgenes (n-Sema5A mice) or littermate controls were trained in contextual fear conditioning, treated with TAM, and then tested for contextual fear 30 days later. In this test, n-Sema5A mice froze at lower levels compared with WT littermate controls (Fig. 5h), indicating that hyper-integration of Sema5A- deficient adult-generated granule cells was sufficient to induce forgetting.

Adult-generated granule cells form input and output synaptic connections, starting around 16 days^17^. Should synaptic integration into hippocampal circuits underlie forgetting, then forgetting should only emerge once Sema5A-deficient adult-generated granule cells have had sufficient time to hyper-integrate (i.e., >16 days). To test this hypothesis, we trained WT and n-Sema5A mice in contextual fear conditioning. Mice were treated with TAM, and then tested 15 days, rather than 30 days, following training. In this test, control and n-Sema5A froze at similar levels (Fig. 5i). These data support the prediction that forgetting should emerge slowly (over weeks) following Sema5A deletion, and the time course of these forgetting effects is consistent with the timing of synaptic integration of adult-generated neurons into hippocampal circuits.

Elevation of hippocampal neurogenesis immediately following training promotes forgetting^10–13^. However, when hippocampal neurogenesis levels are experimentally elevated several weeks following training, forgetting is not observed^12^. The absence of neurogenesis-mediated forgetting at time-points remote to training most likely reflects systems-level reorganization of contextual fear memories^34^. As contextual fear memories age, they become progressively less dependent on the hippocampus^35^, and therefore insensitive to manipulations that impact hippocampal function. Therefore, we predicted that delaying Sema5A deletion until 30 days following training (a time-point when contextual fear memories no longer depend on the hippocampus for expression) would not impact subsequent freezing levels. Accordingly, WT and n-Sema5A were trained in contextual fear conditioning. Thirty days following training, they were treated with TAM, and then tested 30 days later. In this test, WT and n-Sema5A froze at similar levels (Fig. 5j). The lack of effect supports the idea that the hippocampus is no longer playing an essential role in the storage and/or expression of contextual fear memories at time points remote to training. More broadly, the temporal specificity of these results suggest that Sema5A deletion from adult-generated granule cells does not non-specifically reduce the propensity of mice to freeze following fear learning (e.g., by reducing anxiety-like states in n-Sema5A mice). Consistent with this, exploratory behavior in the open field was not altered in n-Sema5A mice (Extended Data Fig. 3e-f).

Deletion of Sema5A from mature, excitatory neurons did affect neuronal morphology (Fig. 6a-f) induce forgetting (Fig. 6g) or alter overall adult neurogenesis levels (Fig. 6h). Similar to our results with Rac1 and Cdh9, these data indicate that the forgetting effects are specific to newborn neurons.

**Figure 6.**
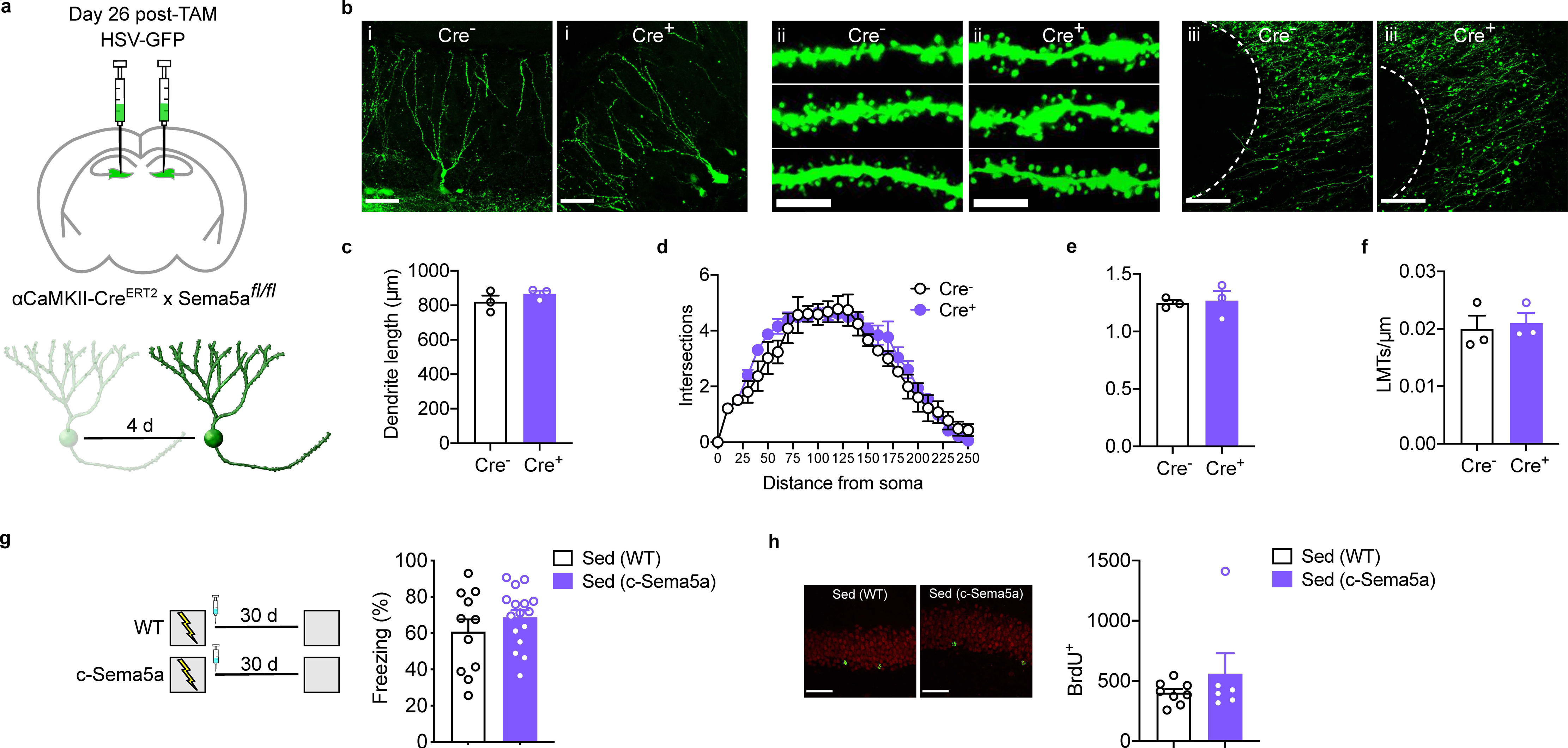
Sema5A deletion in mature neurons does not impact morphology or prevent exercise-induced forgetting. (**a**) Experimental design. c-Sema5A mice were treated with TAM, and then micro-infused with HSV expressing GFP 26 d later. The morphology of infected cells was assessed 4 days later. (**b**) Representative images from hippocampus showing (i) dendritic morphology (scale bar = 50 µm), (ii) dendritic segments (scale bar = 10 µm) and (iii) LMTs (scale bar = 50 µm) of infected cells. Sema5A deletion (Cre^+^ cf. Cre) did not affect (**c**) total dendrite length (*t*_4_ = 1.14, *P* = 0.32), (**d**) dendritic complexity (ANOVA, no main group effect: *F*_1,4_ = 3.43, *P* = 0.14; main distance effect: *F*_27,108_ = 63.05, *P* < 0.0001; no group × distance interaction: *F*_27,108_ = 0.89, *P* = 0.63), (**e**) spine density (*t*_4_ = 0.23, *P* = 0.83) nor (**f**) LMT density (*t*_4_ = 0.35, *P* = 0.74). (**g**) (left) Experimental design. WT and c-Sema5A mice were trained in contextual fear conditioning, and tested 30 d later. One day after training mice were treated with TAM to induce Sema5A deletion in mature neurons. (right) WT and c-Sema5A mice froze at equivalent levels during the memory test (*t*_25_ = 1.09, *P* = 0.29). (**h**) (left) Representative images showing BrdU-labeled cells in DG (scale bar = 50µm). (right) BrdU^+^ cell counts were similar in WT and c-Sema5A mice (*t*_12_ = 1.09, *P* = 0.29).

### Optogenetic activation of adult-generated granule cells promotes dendritic growth and spinogenesis

We asked whether optogenetic activation of young, adult-generated granule cells would accelerate their development and integration. An RV expressing a GFP tagged, excitatory opsin (RV-ChR2-GFP) or control virus (RV-GFP) was injected into the dorsal DG of adult WT mice, and optrodes were implanted bilaterally above the injection site. Starting 2 days later, mice received daily photo-stimulation (3 × 1 min [spaced at 3 min intervals] at 10 Hz) over 15 days.

Fifteen days following completion of photo-stimulation, numerous infected granule cells were identified in the innermost layers of the granule cell layer in the DG (Fig. 7a-b). Similar numbers of control (GFP) and ChR2-expressing infected granule cells were identified (Fig. 7c), suggesting that activation did not alter survival rates of infected adult-generated granule cells (for convergent evidence using BrdU and DCX labeling, see Extended Data Fig. 4). However, ChR2-expressing neurons exhibited increased total dendrite length, dendritic complexity (with increased dendritic branching at higher orders), spine and LMT density (Fig. 7d-g). These results indicate that photoactivation induces hyper-integration of adult-generated granule cells, resulting in increased input and output connectivity. They extend earlier studies showing that chemogenetic activation of adult-generated granule cells accelerates their integration into hippocampal circuits, without affecting survival^36^.

**Figure 7.**
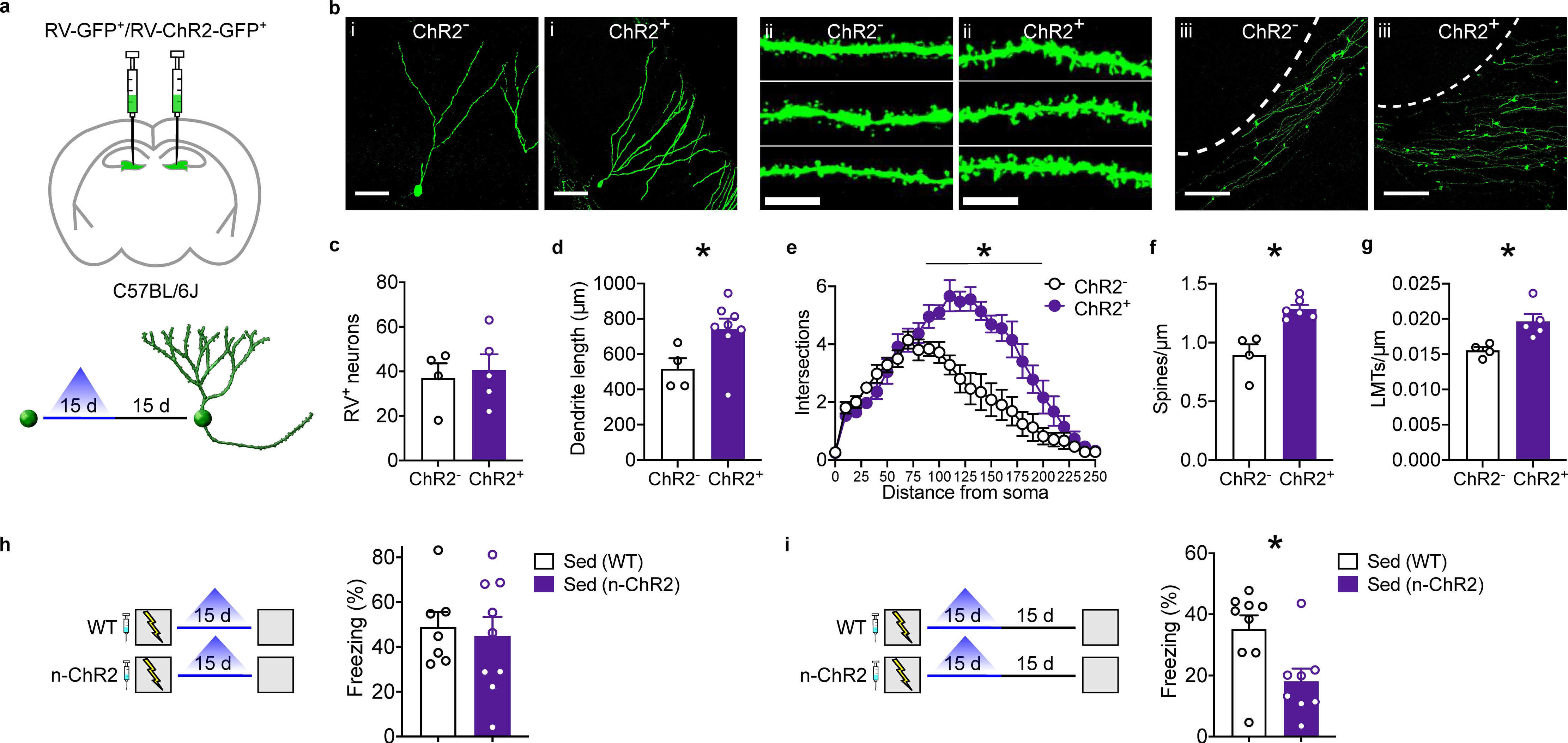
Optogenetic activation of newborn neurons induces hyper-integration and forgetting. (**a**) Experimental design. RV expressing ChR2-GFP or GFP alone was micro- injected into the DG. Blue laser stimulation was delivered daily for 15 d, and morphology of infected adult-generated granule cell assessed 15 d later. (**b**) Representative images from hippocampus showing (i) dendritic morphology (scale bar = 50 µm), (ii) dendritic segments (scale bar = 10 µm) and (iii) LMTs (scale bar = 10 µm) of infected adult-generated granule cells. Photostimulation did not affect (**c**) survival of adult-generated granule cells (*t*_7_ = 0.36, *P* = 0.73), but increased (**d**) total dendrite length (*t*_10_ = 2.35, *P* < 0.05), (**e**) dendritic complexity (ANOVA, main group effect: *F*_1,8_ = 20.47, *P* <0.005; main distance effect: *F*_24,216_ = 31.57, *P* < 0.0001; group × distance interaction: *F*_24,216_ = 6.19, *P* < 0.0001), (**f**) spine density (*t*_8_ = 4.61, *P* < 0.005), and (**g**) LMT density (*t*_7_ = 3.17, *P* < 0.05). (**h**) (left) Experimental design. WT and n-ChR2 mice were treated with TAM, trained in contextual fear conditioning, and tested 15 d later. Optogenetic stimulation was delivered daily for 15 days, starting immediately following training. (right) Optogenetic activation did not induce forgetting at the shorter delay in n-ChR2 mice (*t*_14_ = 0.35, *P* = 0.73). (**i**) (left) Experimental design. WT and n-ChR2 mice were treated with TAM, trained in contextual fear conditioning, and tested 30 d later. Optogenetic stimulation was delivered daily for 15 d, starting one day following training. (right) Optogenetic activation induced forgetting in n-ChR2 mice (*t*_15_ = 2.75, *P* < 0.05).

### Increased integration of ChR2-expressing granule cells induces forgetting

To test whether hyper-integration of adult-generated granule cells following photo-stimulation would be sufficient to induce forgetting, we crossed nestin-Cre^ERT2^ mice with mice that express ChR2 in a Cre-recombinase dependent manner. In adult offspring of this cross expressing both transgenes (n-ChR2 mice), TAM treatment induces excision of a STOP codon and ChR2 expression in nestin^+^ cells and their progeny. WT and n-ChR2 mice were treated with TAM, and then trained in contextual fear conditioning one day later. Mice received 15 days of photo- stimulation starting one day following training, and then were tested either 1 day or 15 days later.

At the shorter retention delay, WT and n-ChR2 mice exhibited equivalent levels of conditioned freezing (Fig. 7h). However, at the longer delay, n-ChR2 mice froze less compared with WT controls (Fig. 7i). The observation that forgetting was only observed at the 30 day time point is consistent with the effects we observed in n-Sema5A mice and with the conclusion that forgetting emerges only once newborn neurons have had sufficient time (i.e., > 16 days) to synaptically integrate into hippocampal circuits.

### Chemogenetic manipulation of adult-generated granule cell activity during consolidation window bidirectionally regulates forgetting

By manipulating how adult-generated neurons integrate into hippocampal circuits, the above experiments suggest that neurogenesis-mediated remodeling of hippocampal circuits represents a continuous and active form of interference. This raises the possibility that eliminating this form of interference, by suppressing the activity of adult-generated neurons during the retention delay, might prevent forgetting induced by post-training exercise. Conversely, promoting this form of interference, by selectively activating adult-generated neurons throughout the retention delay, might be sufficient to induce forgetting in situations where forgetting is not typically observed.

To selectively inhibit newborn neurons during the post-training period, we crossed nestin- Cre^ERT2^ mice with mice that express the inhibitory DREADD, hM4D_i_, in a Cre-recombinase dependent manner. In adult offspring of this cross expressing both transgenes (n-hM4D_i_ mice), TAM treatment induces excision of a STOP codon and hM4D_i_ expression in nestin^+^ cells and their progeny. Adult n-hM4D_i_ mice and their littermate controls were trained in contextual fear conditioning and tested 30 days later. Following training, mice were treated with TAM, and housed either conventionally (‘sedentary’ groups) or with continuous, home cage access to a running wheel (‘exercise’ groups). Throughout this retention delay, all groups of mice were administered the DREADD ligand CNO via their drinking water. We found that exercise-induced forgetting was prevented in mice expressing hM4D_i_ in adult-generated granule cells (Fig. 8a). Additional control groups confirmed that this effect depended on expression of both the DREADD receptor and CNO administration, and that chemogenetic inhibition of newborn neurons did not impact overall levels of neurogenesis (Extended Data Fig. 5a,c).

**Figure 8.**
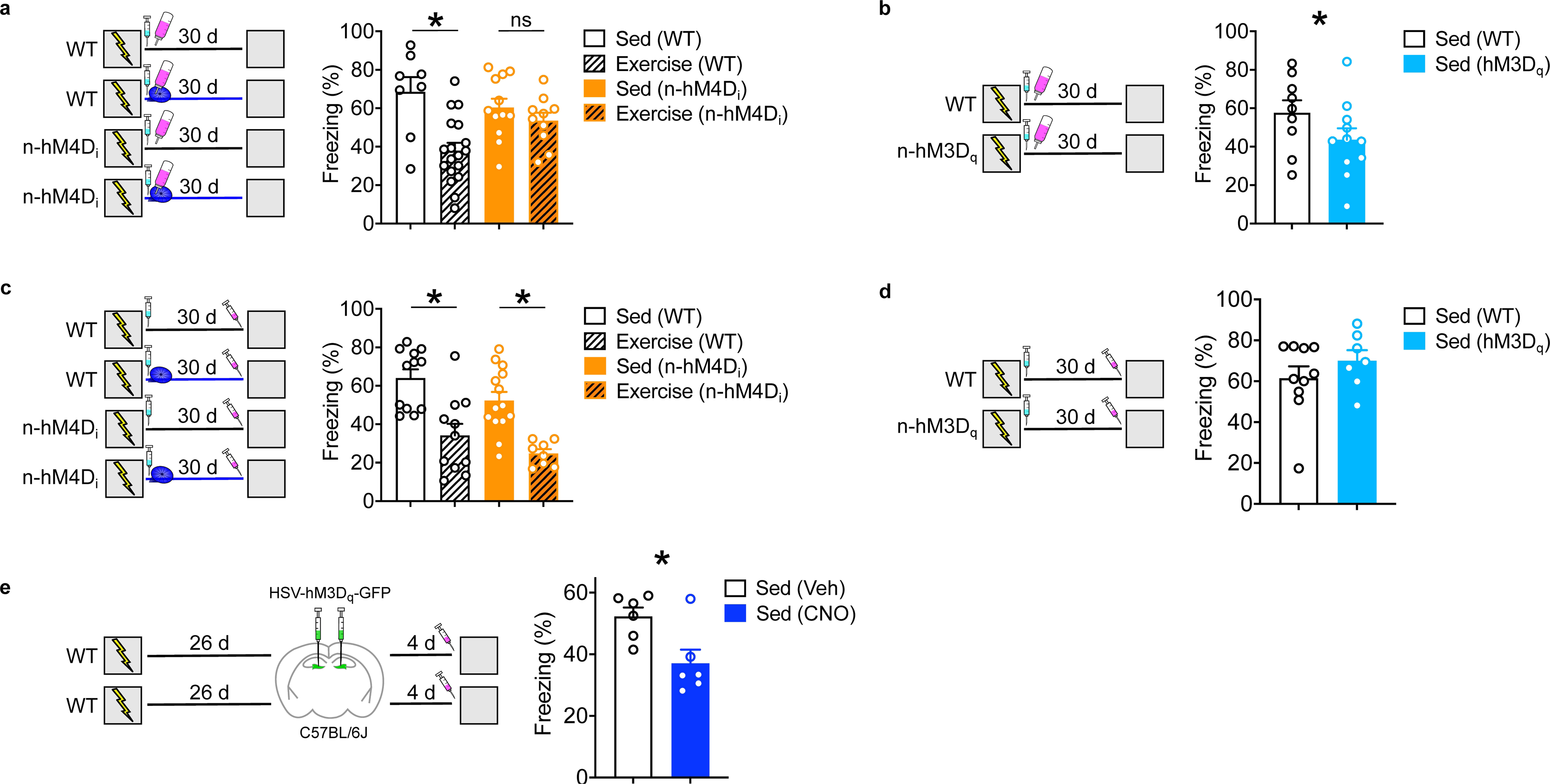
Chemogenetic modulation of newborn neurons regulates forgetting. (**a**) (left) Experimental design. n-hM4D_i_ mice were trained in contextual fear conditioning. Mice either had access to an exercise wheel (‘Exercise’ groups) or were housed conventionally (‘Sed’ groups), and then were tested 30 d later. One day following training, mice were treated with TAM to induce expression of hM4D_i_ in NPCs and their progeny. Mice received CNO (via drinking water) throughout the retention period. (right) Exercise reduced freezing in WT, but not n-hM4D_i_, mice (ANOVA, no main genotype effect: *F*_1,44_ = 0.51, *P* = 0.48; main exercise effect: *F*_1,44_ = 13.39, *P* < 0.001; genotype × exercise interaction: *F*_1,44_ = 5.41, *P* < 0.05). (**b**) (left) Experimental design. n- hM3D_q_ mice were trained in contextual fear conditioning and tested 30 d later. One day following training, mice were treated with TAM to induce expression of hM3D_q_ in NPCs and their progeny. Mice received CNO (via drinking water) throughout the retention period. (right) Freezing was reduced in n-hM3D_q_ compared to WT mice (*t*_18_ = 2.86, *P* < 0.05). (**c**) (left) Experimental design. n-hM4D_i_ mice were trained in contextual fear conditioning. Mice either had access to an exercise wheel (‘Exercise’ groups) or were housed conventionally (‘Sed’ groups), and then were tested 30 d later. One day following training, mice were treated with TAM to induce expression of hM4D_i_ in NPCs and their progeny. Mice received CNO prior to the memory test. (right) Exercise reduced freezing in both WT and n-hM4D_i_ mice (ANOVA, main genotype effect: *F*_1,41_ = 4.67, *P* < 0.05; main exercise effect: *F*_1,41_ = 34.59, *P* < 0.001; no genotype × exercise interaction: *F*_1,41_ = 0.06, *P* = 0.81). (**d**) (left) Experimental design. n-hM3D_q_ mice were trained in contextual fear conditioning and tested 30 d later. One day following training, mice were treated with TAM to induce expression of hM3D_q_ in NPCs and their progeny. Mice received CNO prior to the memory test. (right) Freezing was equivalent in i-hM3D_q_ and WT mice (*t*_12_ = 1.85, *P* = 0.09). (**e**) (left) Experimental design. WT mice were trained in contextual fear conditioning and tested 30 d later. Four d prior to testing, HSV-hM3D_q_ was micro-infused into the DG to infect mature, excitatory neurons. Mice were treated with CNO prior to testing. (right) Freezing was reduced in CNO-treated mice (*t*_10_ = 2.86, *P* < 0.05).

To selectively activate newborn neurons during the post-training period, we crossed nestin- Cre^ERT2^ mice with mice that express the excitatory DREADD, hM3D_q_, in a Cre recombinase- dependent manner. Adult offspring of this cross expressing both transgenes (n-hM3D_q_ mice) and their littermate controls were trained in contextual fear conditioning and tested 30 days later. Following training, the mice were treated with TAM and had continuous access to CNO via their drinking water. In the test, the n-hM3D_q_ experimental mice treated with TAM froze less than their littermate controls, indicating that continuous activation of newborn neurons during the retention delay weakened contextual fear memories (Fig. 8b). Additional control groups confirmed that this effect depended on expression of both DREADD and the presence of CNO, and that chemogenetic activation of newborn neurons did not impact overall levels of neurogenesis (Extended Data Fig. 5b,d).

Similar chemogenetic manipulations of adult-generated neurons at the time of testing (rather than throughout the retention period) had no effect on memory retrieval (Fig. 8c-d). In contrast, chemogenetic activation of mature DG granule cells at the time of testing impaired retrieval of contextual fear memory (Fig. 8e). These results indicate that indiscriminate activation of mature excitatory neurons (but not newborn neurons) interferes with memory retrieval, consistent with recent studies^37–40^.

### Exercise-induced forgetting is associated with reduced retrieval-induced reinstatement of encoding patterns of activation in the hippocampus

Memory retrieval is generally thought to occur when patterns of neural activity that were present during encoding are reinstated^6,41–43^. Given our evidence that newborn neurons promote forgetting by structurally remodelling hippocampal circuity, we predicted that rates of engram reinstatement would be reduced in mice that have undergone neurogenesis-mediated forgetting. Using an ensemble-level engram-tagging approach, we crossed mice in which the TAM-dependent recombinase Cre^ERT2^ is expressed in an activity-dependent manner from the loci of the Arc gene (Arc-Cre^ERT2^, or Arc-TRAP, mice) with mice expressing a floxed-stop tdTomato (tdT) cassette^44,45^. In offspring expressing both transgenes, cells active shortly after TAM injection permanently express a tdT tag. To examine whether engram neurons are reactivated after a memory test, we examined expression of the activity-regulated gene, c-Fos, in the DG, CA3 and CA1 regions of the hippocampus using immunohistochemistry (Fig. 9a).

**Figure 9.**
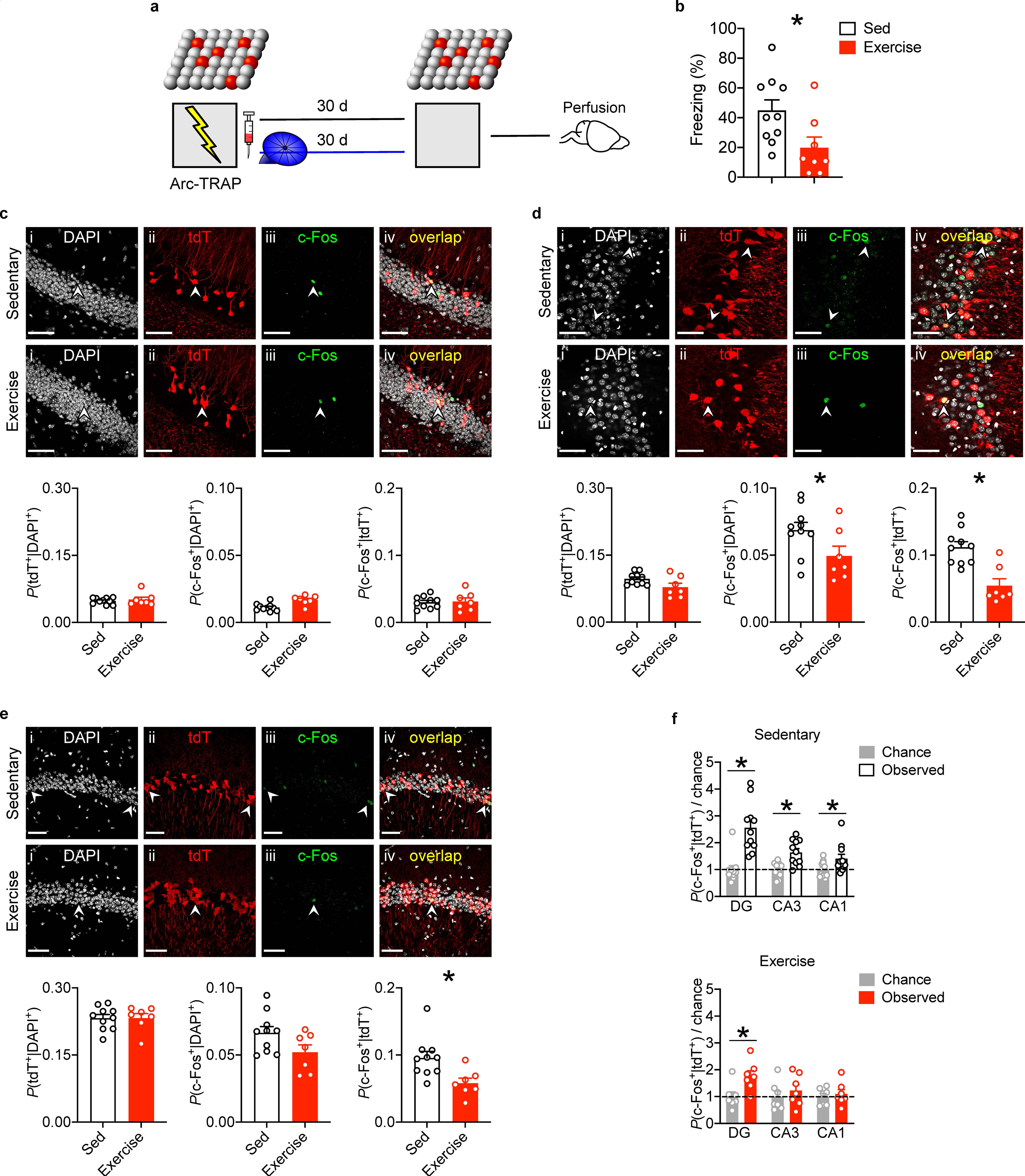
Exercise-induced forgetting reduces engram reactivation. (**a**) Experimental design. Arc-TRAP mice express the TAM-dependent recombinase Cre^ERT2^ in an activity- dependent manner from the loci of the immediate early gene, Arc. Arc-TRAP mice were trained in contextual fear conditioning and tested 30 d later. During then retention delay, mice had home cage access to an exercise wheel (‘Exercise’ group) or were housed conventionally (‘Sedentary’ group). (**b**) Post-training exercise reduced freezing (*t*_16_ = 2.45, *P* < 0.05). (**c**) Representative images of tdT^+^ (tagged encoding ensemble) and c-Fos^+^ (retrieval induced activation) cells in DG. The probability of tdT^+^ neurons (*t*_15_ = 0.56, *P* = 0.59), c-Fos+ neurons (*t*_15_ = 3.62, *P* < 0.005), and reactivation (i.e., *P*(c-Fos^+^│tdT^+^) (*t*_15_ = 0.20, *P* = 0.85) in Sed vs. Exercise mice. (**d**) Representative images of tdT^+^ and c-Fos^+^ cells in CA3. The probability of tdT^+^ neurons (*t*_15_ = 2.08, *P* = 0.055), c-Fos^+^ neurons (*t*_15_ = 2.11, *P* = 0.05) and reactivation (*t*_15_ = 4.34, *P* < 0.005) in Sed vs. Exercise mice. (**e**) Representative images of tdT^+^ and c-Fos^+^ cells in CA1. The probability of tdT^+^ neurons (*t*_15_ = 0.01, *P* = 0.99), c-Fos^+^ neurons (*t*_15_ = 1.99, *P* = 0.064), and reactivation (*t*_15_ = 2.84, *P* < 0.05) in Sed vs. Exercise mice. (**f**) The probability of reactivation relative to chance (i.e., *P*(c-Fos^+^│tdT^+^)/chance) for sedentary mice. Observed reactivation varied by region and exceeded chance in all regions (ANOVA, main region effect: *F*_2,30_ = 4.69, *P* < 0.05; main observed vs chance effect: *F*_1,30_ = 85.72, *P* < 0.0001; region × observed/chance interaction: *F*_2,30_ = 14.05, *P* < 0.0001; LSD post-hocs: DG [*P* < 0.001], CA3 [*P* < 0.001], CA1 [*P* < 0.05]). (**g**) The probability of reactivation relative to chance (i.e., *P*(c- Fos^+^│tdT^+^)/chance) for exercise mice (ANOVA, main region effect: *F*_2,18_ = 2.31, *P* = 0.13; main observed vs chance effect: *F*_1,18_ = 4.77, *P* < 0.05; region × observed/chance interaction: *F*_1,18_ = 1.49, *P* = 0.25). Reactivation exceeded chance in DG [*P* < 0.05], but not CA3 [*P* = 0.43] or CA1 [*P* = 0.74]). Scale bars = 50 µm.

Arc-TRAP-tdT mice were trained in contextual fear conditioning, and injected with TAM immediately after training to tag encoding ensembles. Mice were returned to their home cages, where they had continuous access to an exercise wheel or were housed conventionally. In a test 30 d later, freezing was reduced in the exercise group (Fig. 9b). The probability of tdT^+^ neurons (i.e., reflecting training activation) was similar in sedentary vs. exercise groups in the DG, CA3 and CA1 (Fig. 9c-e). The probability of c-Fos^+^ neurons (i.e., reflecting test activation) was similar in sedentary vs. exercise groups in the DG and CA1 (Fig. 9a), but was reduced in the exercise group in CA3 (Fig. 9d). Most importantly, the probability of neurons tagged during training being reactivated during testing (i.e., *P*(tdT^+^│c-Fos^+^)) was reduced in CA3 and CA1 regions in the exercise group (Fig. 9d-e). To directly compare reactivation rates across regions and groups, we compared observed reactivation rates to those expected by chance. In sedentary mice, these analyses revealed above chance levels of reactivation in all three subregions of the hippocampus analyzed (Fig. 9f). In contrast, reactivation rates were only above chance levels in the DG in mice in the exercise group, and did not differ from chance in CA3 and CA1 (Fig. 9g).

### Reactivation of DG encoding ensembles leads to memory recovery following forgetting

As noted above, reinstatement levels remained above chance in the DG in the exercise group. This indicates that neural reinstatement is not completely abolished following post-training exercise, and raises the possibility that artificial engram reactivation during a retrieval test might recruit residual engram circuitry and restore freezing levels to those observed in control mice.

To test this possibility, we used a second ensemble tagging approach (robust activity marking; RAM) to express the excitatory DREADD hM3D_q_ in DG neurons activated during initial learning^46^. This tet-OFF based tagging approach labels active ensembles when doxycycline (DOX) is removed from the diet. Accordingly, DOX-fed mice were micro-infused with AAV-RAM- hM3Dq-mCherry. Three weeks later DOX was removed from their diet, and they were trained in contextual fear conditioning. Upon return to their home cage, mice resumed their DOX diet, and either had access to an exercise wheel or were housed conventionally. Thirty days later, mice were tested. During this test, mice were treated with CNO (or vehicle) in order to reactivate DG granule cells that were previously active during encoding (Fig. 10a).

**Figure 10.**
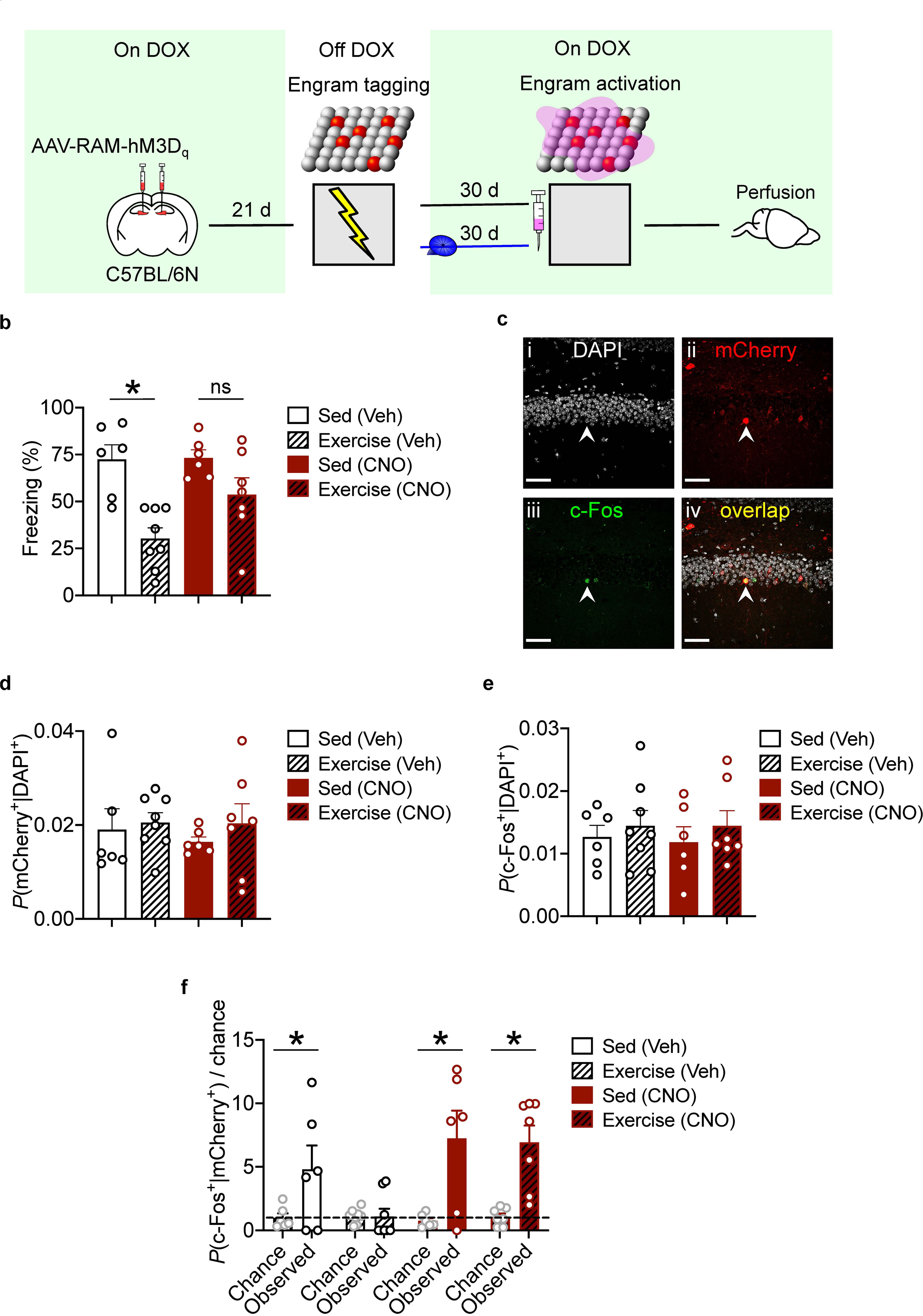
Recovery of fear memory via reactivating of DG ensembles that were active during encoding. (**a**) Experimental design. Mice were micro-infused with AAV-RAM-hM3D_q_- mCherry and trained in contextual fear conditioning in the absence of DOX in order to tag active neurons. Mice either had access to an exercise wheel (‘Exercise’ groups) or were housed conventionally (‘Sed’ groups), and then were tested 30 d later. Prior to testing, mice were treated with CNO (or Veh) in order to activate DG ensembles that were active during initial learning. (**b**) Post-training exercise reduced freezing in vehicle-treated mice (planned t-test: *t*_12_ = 4.52, *P* < 0.001). Reactivation of tagged engram cells in CNO treated mice restored freezing to control levels (planned t-tests comparing exercise [CNO] vs. sedentary [CNO] mice [*t*_12_ = 4.52, *P* = 0.09], exercise [CNO] vs. exercise [Veh] [*t*_13_ = 2.29, *P* < 0.05], and exercise [CNO] vs. sedentary [Veh] [*t*_12_ = 1.35, *P* = 0.21]. **(c)** Representative images from DG with mCherry^+^(encoding ensemble), c-Fos^+^ (retrieval ensemble) and overlap (encoding neurons that were reactivated by retrieval). (**d**) mCherry^+^ neurons (i.e., encoding ensemble) did not differ among groups (ANOVA, no main exercise effect: *F*_1,23_ = 0.69, *P* = 0.41; no main drug effect: *F*_1,23_ = 0.18, *P* = 0.67; no genotype × exercise interaction: *F*_1,23_ = 0.14, *P* = 0.71). (**e**) c-Fos^+^ (i.e., retrieval ensembles) did not differ among groups (ANOVA, no main exercise effect: *F*_1,23_ = 0.86, *P* = 0.36; no main drug effect: *F*_1,23_ = 0.03, *P* = 0.86; no genotype × exercise interaction: *F*_1,23_ = 0.03, *P* = 0.86) (**f**) reactivation relative to chance (*P*[c-Fos^+^│mCherry^+^]/chance) differed across groups (ANOVA, main group effect *F*_3,23_ = 3.52, *P* < 0.05; main observed vs. chance effect: *F*_1,23_ = 28.84, *P* < 0.001; group × observed vs. chance interaction: *F*_3,23_ = 4.22, *P* < 0.001). Reactivation exceeded chance in Sed (Veh) [*P* < 0.05], Sed (CNO) [*P* < 0.001] and Exercise (CNO) [*P* < 0.001], but not Exercise (Veh) [*P* = 0.43] groups. Scale bars = 50 µm.

In the test, post-training exercise reduced freezing levels in vehicle-treated mice (Fig. 10b), as expected. However, rates of forgetting were significantly attenuated in CNO-treated mice, indicating that chemogenetic activation of encoding ensembles of DG granule cells is sufficient to restore forgotten contextual fear memories. Following testing, we additionally performed immunohistochemistry for c-Fos in order to assess the extent to which encoding ensembles were reactivated during testing in the various groups (Fig. 10c). The size of the encoding (i.e., mCherry^+^) and retrieval (i.e., c-Fos^+^) ensembles were similar across groups (Fig. 10d-e).

However, reactivation rates differed (Fig. 10f). In vehicle-treated mice, reactivation rates were reduced in mice that exercised following training, consistent with their reduced freezing levels. In CNO-treated mice, reactivation rates were similar in sedentary and exercise groups, and did not differ from mice in the vehicle-treated, sedentary group. These results indicate that forgetting is associated with reduced engram reinstatement and that reactivating tagged engram cells in the DG is sufficient to reverse forgetting, consistent with previous studies^47–49^.

## DISCUSSION

In adult rodents, post-training increases in hippocampal neurogenesis induce forgetting of hippocampus-dependent memories. Using both gain and loss of function approaches, here we tested whether neurogenesis-mediated forgetting is caused by changes in connectivity. We designed a variety of NPC-specific interventions that alter how new neurons integrate into hippocampal circuits. One class of intervention (deletion of Sema5A or optogenetic activation) promoted hyper-integration, with new neurons exhibiting increased input (e.g., increased dendritic complexity and spine density) and output connectivity (e.g., increased LMT density). We found that inducing hyper-integration of new neurons following training was sufficient to induce forgetting of contextual fear memories. A similar pattern of effects was observed when we chemogenetically activated newborn neurons throughout the retention delay. Another class of interventions (deletion of Rac1 or Cdh9) promoted hypo-integration of new neurons into hippocampal circuits, with new neurons exhibiting decreased input (e.g., decreased dendritic complexity and spine density) and/or output connectivity (e.g., decreased LMT density). We found that inducing hypo-integration of new neurons following training prevented exercise- induced forgetting of contextual fear memories. A similar pattern of effects was observed when we chemogenetically inhibited newborn neurons throughout the retention delay. Because these interventions did not affect survival of newborn neurons, these experiments indicate that changes in patterns of connectivity are necessary and sufficient for neurogenesis-mediated forgetting. Moreover, they suggest that neurogenesis-mediated remodeling of hippocampal circuits represents a continuous and active form of interference that gradually reduces accessibility of engrams underlying hippocampal memories. Consistent with this, using engram- labeling approaches, we found that exercise-induced forgetting was associated with reduced engram reactivation.

Several studies have shown that interventions that increase hippocampal neurogenesis induce forgetting in adult rodents^10–14^. However, in many cases, these interventions lack specificity (e.g., exercise, systemic treatment with the NMDA antagonist memantine). Since they additionally affect processes other than hippocampal neurogenesis, it is not possible to exclude the possibility that non-neurogenic mechanisms contribute to forgetting. The interventions in the current study targeted newborn neurons, and therefore directly link adult neurogenesis to forgetting. Our findings additionally suggest that forgetting depends on synaptic integration of newborn neurons into hippocampal circuits. After promoting hyper-integration of newborn neurons, memory was unaffected when tested 15 days later. However, forgetting was apparent at the 30 day retention delay. The gradual emergence of forgetting matches the time course of synaptic integration of new neurons. As new neurons migrate into the granule cell layer they begin to extend an axon towards CA3 and dendrites towards the molecular layer, forming input and output synapses after ∼16 days^17,19,20,50,51^.

The effects of our manipulations were temporally graded, affecting recent, but not remote, memories. The circuits supporting contextual fear memories reorganize over time in a process known as systems consolidation^34^. Whereas days following training, retrieval of contextual fear memories depends on the hippocampus, weeks following training these memories may be expressed independently of the hippocampus. Consistent with this time course, promoting hyper-integration of newborn neurons only produced forgetting when induced one day, but not month, following training. Similarly, promoting hypo-integration of newborn neurons only prevented forgetting when induced following training, and not at more remote time points. Post- training exercise produces similar temporally-graded forgetting effects^12^, and therefore suggest that hippocampus-dependent memories are only transiently sensitive to neurogenesis-mediated remodeling of hippocampal circuits. Their vulnerability decreases over weeks, presumably as these memories become fully consolidated in the cortex.

Forgetting was only affected when our manipulations targeted newborn neurons. When gene deletions were targeted to excitatory, mature neurons throughout the brain (i.e., αCaMKII^+^ cells) we found no effects on forgetting. Whereas conditional deletion of Rac1 or Cdh9 from nestin^+^ cells blocked exercise-induced forgetting, conditional deletion of Rac1 or Cdh9 from αCaMKII^+^ cells had no effect. Likewise, whereas conditional deletion of Sema5A from nestin^+^ cells induced forgetting, conditional deletion of Sema5A from αCaMKII^+^ cells had no effect. This pattern of results is striking given that mature excitatory neurons outnumber newborn neurons by several orders of magnitude. This may indicate that thresholds for inducing structural changes differ in mature compared to newborn neurons. Indeed, deletion of Rac1, Cdh9 and Sema5A did not impact the morphology of mature, excitatory neurons, including dendritic complexity, spine and LMT density. In contrast, equivalent manipulations in newborn neurons caused more pronounced morphological changes, impacting the formation of input and output connections.

To our knowledge, the effects of manipulating Sema5A or Cdh9 on memory stability have not previously been examined. However, the impact of manipulating Rac1 in mature, excitatory neurons has been examined previously^52^. In this study, spatial memory retention was examined in conventionally-housed mice lacking Rac1 in forebrain, excitatory (αCaMKII^+^) neurons. Similar to our results using contextual fear, these Rac1 mutants exhibited normal expression of water maze memories up to 5 weeks following training. In contrast, over-expression of a constitutively active form of Rac1 in excitatory forebrain neurons accelerates LTP decay and forgetting of object memories^53^.

How does neurogenesis-mediated circuit remodeling induce forgetting? Memory formation involves the strengthening of synaptic connections among neurons that are co-active during encoding^42,43,54^, and subsequent reactivation of these neuronal ensembles (or engrams) then leads to successful memory retrieval^41^. As new neurons integrate into hippocampal circuits, they compete with existing neurons for input and output connections^18–20^. Therefore, as these connectivity changes accumulate, the likelihood that a given input (e.g., retrieval cue) faithfully reactivates an engram may fade^22–24,55,56^.

We found evidence for reduced engram reactivation following exercise-induced forgetting of contextual fear memories. During the memory test, reactivation rates in the exercise group were at chance levels in CA1 and CA3. However, reactivation rates, while reduced, remained above chance levels in the DG. The fact that engram reactivation was not reduced to chance levels in DG raises two interrelated issues.

First, this pattern suggests that the locus of neurogenesis-mediated forgetting is at the mossy fiber-CA3 synapse (cf. the EC-DG performant path synapse). This conclusion is consistent with earlier computational modeling which explored the impact of middle layer neurogenesis on forgetting in a categorization task^23^. Across a broad parameter space, these *in silico* experiments indicated that memory stability was more sensitive to changes in output, rather than input, adult-generated granule cell connectivity. Analysis of the n-Cdh9 mice also support this conclusion. In these mice, morphological changes were restricted to output connections (i.e., reduced LMT size and density), with no changes in input connectivity (i.e., dendritic length, dendritic complexity, or spine density). Importantly, induction of this partial hypo-integration phenotype post-training prevented exercise-induced forgetting, suggesting that neurogenesis- induced remodeling of mossy fiber-CA3 synapses regulates forgetting. Given that overall retrieval-induced activity was reduced in CA3 in the mice that exercised, one possibility is that circuit remodeling weakens engram (DG)◊ engram (CA3) connectivity, resulting in inefficient CA3 pattern completion and subsequent retrieval failure. Indeed, other studies have similarly shown that retrieval success correlates with reactivation of CA3, but not DG, engram neurons^57,58^.

An alternative possibility is that neurogenesis-dependent remodeling induces changes in excitatory-inhibitory balance that lead to forgetting. Although not characterized here, newborn granule cells also form connections with local inhibitory circuits in the hippocampus, including parvalbumin^+^ inhibitory interneurons in stratum lucidum which provide feedforward inhibition onto principal neurons in CA3^24,51,59–61^. Therefore, increased newborn neuron-inhibitory interneuron connectivity might promote increased feedforward inhibition, resulting in blunted activation of CA3 engram cells and retrieval failure^60^. However, our finding that transient chemogenetic activation of newborn neurons during a retrieval test did not affect contextual fear memory retrieval suggests this possibility is less likely.

Second, post-training exercise did not reduce reactivation rates to chance levels in the DG, indicating that forgetting is incomplete. This suggests that neurogenesis-mediated remodeling of hippocampal circuits does not completely overwrite memories, and that residual ‘engram’ circuitry remains. Indeed, while presentation of natural retrieval cues was associated with reduced freezing and chance levels of engram reactivation in CA3/CA1, it is possible that boosting activation of DG engram cells in the presence of natural cues would lead to more efficient engram reactivation rates in these downstream regions. Consistent with this prediction, we found that chemogenetic activation of DG engram cells during a memory test alleviated forgetting.

More broadly, this form of neurogenesis-mediated forgetting represents one of several ways in which the brain forgets^6,9^. Recent studies have identified several putative forgetting mechanisms. In addition to neurogenesis-mediated remodeling of hippocampal circuits, these include microglia-dependent synapse elimination^15^, synaptic weakening via AMPA receptor endocytosis^62,63^, and spine instability/turnover^9,64^. While these may differ in terms of mechanism and time course, nonetheless they appear to converge on the common principle that circuit stability is required for memory stability, and that mechanisms that disrupt circuit architecture promote forgetting^65^. These forms of natural forgetting may represent adaptive solutions to avoid over-fitting to specific past instances, allowing organisms to better generalize to future events^6,66^.

## METHODS and METHODS

### Mice

Procedures were approved by Hospital for Sick Children Animal Care and Use Committee and conducted in accordance with CCAC and US National Institutes of Health (NIH) guidelines. Mice were bred and maintained in the Laboratory Animals Services vivarium at The Hospital for Sick Children. After weaning at post-natal day 21 (P21), same sex mice were group-housed in standard mouse housing cages (2–5 per cage) and maintained on a 12 h light/dark cycle.

Behavioral testing occurred during the light phase. Roughly equal numbers of male and female mice were used in all experiments. Behavioral and morphological experiments were conducted by experimenters unaware of mouse genotype and treatment of the mouse. All mice were maintained on a C57BL/6J background except for the ChR2 line which were on maintained on a 129/SvEv background.

Briefly, we used mouse lines floxed for our genes of interest and used either viral or transgenic methods of delivering Cre recombinase to knock out genes or express ChR2, DREADDs, or fluorescent proteins in particular cell types at particular times. In most experiments, we used a Cre recombinase line that expresses Cre in a TAM-dependent manner in NPCs (nestin-Cre^ERT2-/+^). We also used a mouse line expressing Cre recombinase in a TAM-dependent manner in excitatory forebrain neurons (CaMKII-Cre^ERT2-/+^) in order to examine the impact of gene deletions in mature neurons in adult mice.

Cdh9 conditional mutant mice were produced by flanking exon 5 of the cdh9 gene with LoxP sites. The targeting vector was generated from Knockout Mouse Project (KOMP) clone 119231, modified by removal of 5’ sequences as described previously^67^. Following electroporation into ES cells, clones were selected with G418 and screened for proper integration. Chimeras were generated by blastocyst injection at the Harvard University Genome Modification Facility and germline transmission was obtained from two independent clones. Offspring were bred to animals expressing FLP recombinase from the beta-actin promoter^68^ to remove the neomycin and an extraneous LoxP site, generating the conditional allele.

<u>For behavioral experiments</u>, we crossed mouse lines floxed for our gene of interest with mice expressing a tamoxifen (TAM)-inducible Cre recombinase in NPCs. We used Nestin-Cre^ERT2-/+^ (line 4 from Imayoshi et al., 2008 [REF: ^69^]) to delete genes of interest from Rac1^fl/fl^ mice (RRID:IMSR_JAX:005550), Cdh9^fl/fl^ mice, and Sema5a^fl/fl^ mice^33^ (generous gift from Dr. Roman Giger, University of Michigan). We refer to the double mutant offspring resulting from this cross as n-Rac1, n-Cdh9 and n-Sema5a mice, respectively. Littermate control mice (i.e., Cre^-^) were used for all experiments. TAM was injected for 5 consecutive d. In most experiments, this started one d after contextual fear conditioning and forgetting examined 15-30 d later. In some experiments, TAM treatment occurred 30 d after conditioning, and forgetting was examined 30 d later.

<u>To control for effects of deleting our genes of interest from developmentally-generated neurons</u> <u>in behavioral experiments</u>, we crossed our floxed lines of mice (Rac1^fl/fl^, Cdh9^fl/fl^ and Sema5a^fl/fl^ mice) with a TAM-dependent Cre recombinase driver line which expresses Cre in adult excitatory forebrain neurons (αCamKII-Cre^ERT2-/+^; RRID:IMSR_JAX:012362). We refer to the double mutant offspring resulting from this cross as c-Rac1, c-Cdh9 and c-Sema5a mice, respectively. Littermate control mice (i.e., Cre^-^) were used for all experiments. As above, TAM was injected for 5 consecutive d after contextual fear conditioning and forgetting examining 30 d later.

<u>To characterize the morphology of adult-generated neurons</u>, we micro-infused retrovirus (RV) expressing GFP and Cre-recombinase (RV-Cre^+^) or GFP alone (RV-Cre^-^) into the DG of mouse lines floxed for our genes of interest (Rac1^fl/fl^, Cdh9^fl/fl^ and Sema5a^fl/fl^ mice). Briefly, 0.5 µl RV expressing either GFP or GFP-IRES-Cre driven by a CAG promoter was micro-infused bilaterally into the dorsal DG of mice. RV-Cre^-^ labels dividing NPCs while RV-Cre^+^ both labels and deletes the gene of interest from dividing NPCs in floxed mice. We examined the morphology of infected neurons 30 d post-RV micro-infusion.

<u>To control for effects of deleting our genes of interest from developmentally-generated neurons</u> <u>in morphological experiments</u>, we used a virus which infects adult excitatory neurons, rather than NPCs. We micro-infused HSV-GFP into αCamKII-Cre^ERT2-/+^ × floxed gene of interest line crosses (c-Rac1, c-Cdh9 and c-Sema5a mice). HSV-GFP randomly infects a sparse population of excitatory neurons^70,71^ which allowed us to analyze the effects of deleting these genes in developmentally-born neurons in adult mice.

<u>To examine the morphological effects of optogenetic activation newborn neurons in wild-type</u> <u>mice</u>, we micro-infused RV expressing ChR2-GFP or GFP alone into wild-type mice (C57BL/6N mice, Taconic Bioscience). ChR2-expressing neurons were stimulated with blue light (see below for details) for 14 d and morphology assessed 16 d later (Day 30).

To examine the behavioral effects of artificially manipulating (via optogenetics and chemogenetics) newborn neurons in wild-type mice, we crossed Nestin-Cre^ERT2-/+^ with mice that express a loxP-flanked STOP cassette upstream of the ChR2-EYFP fusion gene at the Rosa 26 locus mice (RRID:IMSR JAX:012569) (n-ChR2 mice). TAM was injected for 3 consecutive d, and then mice were contextually fear conditioned. ChR2-expressing neurons were optogenetically stimulated with blue light for 14 d (details below), and forgetting assessed either 1 or 15 d later. In chemogenetic experiments, we crossed Nestin Cre^ERT2-/+^ with mice that express a loxP-flanked STOP cassette upstream of hM3D_q_-mCitrine or hM4D_i_-mCitrine fusion genes at the Rosa 26 locus (RRID:IMSR JAX:026220 and RRID:IMSR JAX:026219) to generate n-hM3D_q_ and n-hM4D_i_ mice, respectively. Following contextual fear conditioning, TAM was injected for 5 consecutive d. Clozapine-N-oxide (CNO; a ligand for selectively activating DREADDs, including hM3D_q_ and hM4D_i_) was administrated either throughout the 30 d memory consolidation period, or 1 h before memory retrieval (details below).

To control for effects of chemogenetically increasing the activity of developmentally-generated neurons in adult wild-type mice on memory retrieval, we injected a HSV-hM3D_q_-GFP into wild- type mice 26 d after contextual fear conditioning. Four d later, memory was tested, 1 h after CNO administration.

<u>To label active neurons</u>, we crossed Arc-Cre^ERT2+/+^ transgenic mice (Arc-TRAP mice, RRID:IMSR_JAX:021881) with mice expressing a floxed-stop tdTomato (tdT) cassette (RRID:IMSR_JAX:007914). In offspring expressing both transgenes, cells active in a small window following 4-OHT injection permanently express tdT.

<u>To chemogenetically activate labelled active neurons</u>, we used a viral vector expressing the robust <u>a</u>ctivity <u>m</u>arking (RAM) to tag active neurons in a temporally-specific (doxycycline [DOX] off) manner^72^. AAV-RAM-hM3D_q_-mCherry was micro-infused into the DG of wild-type mice. Mice were maintained on a DOX diet and removed from this diet 48 h before contextual fear conditioning, such that neurons active during conditioning would be tagged and express the chemogenetic actuator hM3D_q_. Mice were returned to a DOX diet 1.5 h following training (details below). To chemogenetically activate tagged engram neurons, CNO (or vehicle) was injected 1 h before the 30 d memory test.

### Surgery

Mice were pre-treated with atropine sulfate (0.1 mg/kg, i.p.), anesthetized with chloral hydrate (400 mg/kg, i.p.), administered meloxicam (2 mg/kg, s.c.) for analgesia, and placed into stereotaxic frames (ASI Instruments). The scalp was incised and retracted, and holes were drilled into the skull above the dorsal DG. Viruses were injected bilaterally via a glass micropipette (for RV, 0.5 µl at a rate of 0.1 µl/min; coordinates: AP −1.9 mm, ML ± 1.5 mm, DV −2.0 mm; for HSV-GFP, 0.1µl (diluted x20) at a rate of 0.1 µl/min; coordinates: AP −1.9mm, ML ± 1.5 mm, DV −2.0 mm; for HSV-hM3D_q_-GFP, 1.5 µl at a rate of 0.1 µl/min; coordinates: AP −1.9mm, ML ± 1.5 mm, DV −2.0 mm; for AAV-RAM-hM3D_q_-mCherry, 1.5 µl at a rate of 0.1 µl/min; coordinates: AP −1.9mm, ML ± 1.5 mm, DV −2.0 mm). Micropipettes were left in place at least 5 min after micro-infusion. The scalp was stitched and Polysporin applied to the wound.

For optogenetic stimulation experiments, optic fibers were implanted above the dorsal DG (AP −1.9 mm, ML ± 1.5 mm, DV −1.7mm). Optic fibers were constructed in-house by attaching a 10 mm piece of 200 mm, optical fiber (with a 0.37 numerical-aperture, NA) to a 1.25 mm zirconia ferrule. Fibers were attached with epoxy resin into ferrules, cut and polished. Optical fibers were stabilized to the skull with screws and dental cement. Dental cement was painted black to minimize light leakage. After surgery, mice were administered 1 ml of 0.9% saline (s.c.) and placed on heated bedding for the duration of the post-surgery recovery period.

### Drugs

#### Tamoxifen

To induce recombination in our Cre-dependent mouse lines, TAM (Sigma, T5648) was injected starting 24-48 h after fear conditioning (170 mg/kg, i.p. for 5 d)^73^. Sugar water was made available to promote hydration on the days of TAM administration. TAM was dissolved in a 10% ethanol/90% sunflower seed solution. Specifically, 30 mg aliquots of TAM were dissolved in 100 ml ethanol by vortexing for ∼30 s. 900 ml sunflower seed oil was added and the solution was vortexed and sonicated until TAM was fully dissolved.

### 4-Hydroxytamoxifen (4-OHT)

Immediately upon removal from the fear conditioning chambers, recombination was induced in Arc-TRAP mice via injection of 4-OHT (Toronto Research Chemicals, Cat no. T006000) (33 mg/kg, i.p. injected at 5 μl/g mouse). 4-OHT was first mixed with 100% ethanol (15 mg/375 μl) and vortexed vigorously. The solution was then placed into a 50 °C chamber, vortexing every 12 min for ∼2 h until fully dissolved. An equal part cremophor (375 μl) was added to create a stock solution that was stored at −20 °C until required. On the day of the experiment, stock solution was mixed at a 1:2 ratio with PBS.

#### Bromodeoxyuridine (BrdU)

To tag dividing cells, BrdU (Sigma, 19-160), a thymidine analog, was co-injected (100 mg/kg, i.p. injected at 10 µl/g mouse) on the final day of TAM administration. BrdU was dissolved in a 50-55 °C PBS solution. This solution was vortexed until BrdU was fully dissolved. BrdU was allowed to cool to room temperature before injecting^74^.

#### Clozapine-N-oxide (CNO)

CNO (Toronto Research Chemicals, C587520) was either administered in drinking water throughout the memory consolidation period (and replaced with normal water 24 h before memory retrieval test) or via i.p. injection 1 h before memory retrieval test. CNO was made in a stock solution of 10 mg/ml in DMSO. Each water bottle received 600 µl CNO stock solution (6 mg CNO/600 µl DMSO) + 150 ml water + 1.5 g sucrose. Mice had no other source of water during this time. CNO was delivered systemically at a dose of 1.0 mg/kg, i.p. (at 10 µl/g mouse).

#### Viruses

All viruses were made in-house.

#### RV

To target NPCs, we used RVs (based on the Moloney murine leukemia virus). These viruses only express in dividing cells^17,75,76^. We used a CAG promoter-driven GFP (Addgene #16664), GFP-IRES-Cre (Addgene #48201), or ChR2-GFP (Addgene #114367). Viral particles were generated as previously described^51,77^. Briefly, retroviruses were prepared by transfecting Platinum-gp cells with plasmids containing an amphotropic envelope (vesicular stomatitis virus- glycoprotein) and the transgenes. Platinum-E cells were then infected to generate a stable virus-producing cell line, and concentrated virus solution was obtained by ultraspeed centrifugation with final titers of ∼1.7 x 10^2^ - 3.5 x 10^9^ units/ml.

#### HSV

To target developmentally-generated neurons in adult mice, we used a replication- defective HSV-GFP: HSV-p1005 (Russo et al., 2009). Transgene expression was driven by the IE 4/5 promoter ^78^. When microinjected into the forebrain regions such as the lateral amygdala or hippocampus, HSV randomly infects approximately 10∼20 % of principal (excitatory) neurons, rather than interneurons^79^, and reaches maximal expression levels 2∼4 d post- injection^80,81^. An hM3D_q_ construct, kindly provided by Dr. Bryan Roth (University of North Carolina), was subcloned into the HSV-p1005 [REF: ^82^] to create the HSV-hM3D_q_-GFP.

#### AAV

To tag active neurons active during contextual fear conditioning, we used the neuronal activity-dependent tagging system, <u>R</u>obust <u>A</u>ctivity <u>M</u>arking (RAM)^72^. The RAM AAV viral vector system tags (with a transgene or fluorophore) active neurons (via a synthetic activity-regulated promoter, P_RAM_, made up of minimal AP-1, Fos and Npas4 promoter sequences) in a temporally-specific fashion (via a doxycycline (DOX) -dependent modified Tet-Off system). pAAV-RAM-d2TTA::TRE-EGFP-WPREpA was a gift from Yingxi Lin (Addgene #84469). We subcloned in the hM3D_q_ construct into this backbone. DOX, administered in the food (40 mg/kg before training and 200 mg/kg after training), prevented hM3D_q_-mCherry expression. The “tagging window” was opened by the withdrawal of DOX-containing food for 48h before training such that active neurons expressing this AAV-RAM-hM3D_q_-mCherry subsequently expressed hM3D_q_-mCherry. Mice were returned to a DOX diet 1.5 h following training.

### Optogenetic stimulation

To characterize the effect of chronic optogenetic activation on the morphological development of adult-generated neurons, RVs expressing GFP (CAG-GFP) or ChR2-GFP (CAG-ChR2-GFP) were injected into the DG of wild-type mice, with optogenetic stimulation beginning 2 d later. To determine the effect of chronic optogenetic stimulation on rates of forgetting, TAM was injected for 3 consecutive d in n-ChR2 mice. Mice were trained in contextual fear conditioning, and optogenetic stimulation began 24 h later. In both experiments, we stimulated the tagged population of newly generated neurons once per d for 14 d. Mice were placed into a Plexiglas container with clean bedding, located in a neutral, novel room. The implanted optic fibers were tethered to a laser source (Laserglow) and controlled by a function generator (Agilent Technologies). Mice received 3 epochs of laser blue light stimulation (473 nm, 10 Hz, 15 ms pulses, 5 vpp, 30% duty cycle, at 0.4 mW from 60-120 s, 300-360 s, and 540-600 s). Mice remained in the Plexiglas container for an additional 60 s and were returned to their home-cage.

### Behavior

#### Contextual fear conditioning and testing

Mice were placed into a fear conditioning chamber (31 cm × 24 cm × 21 cm; Med Associates) with shock-grid floors (bars 3.2 mm diameter, spaced 7.9 mm apart). Footshocks (0.5 mA, 2 s duration) were delivered 120 s, 180 s, and 240 s after placement in the chamber. Mice were removed from the conditioning chamber 60 s after the final shock and returned to their home cage. Either 15, 30, or 60 d later, mice were placed back into the contextual fear chamber for a 5-min memory test. The amount of time mice spent freezing (percentage freezing, with minimum bout of 1 s) was monitored with overhead cameras and calculated using automatic scoring software FreezeFrame v.3.32 (Actimetrics). Freezing is an active defensive response defined as cessation of movement, except for breathing^83,84^.

#### Running wheels

To increase rates of hippocampal neurogenesis post-conditioning, running wheels were placed in the home cage ∼3 h after contextual fear training^10^. To promote acclimation, the running wheel was rubbed with bedding from the home-cage of the mice. To facilitate its use, mice were placed on the running wheel for ∼30 s each for 2 d after the wheels were put in the home cage. Running wheels remained in the cage until the end of the experiment.

#### Open field

To determine whether deletion of Rac1, Cdh9, or Sema5a from adult-generated neurons alters anxiety-like behavior, we tested n-Rac1, n-Cdh9, and n-Sema5a mice in an open field task. Mice were injected with TAM for 5 consecutive d (as in above experiments), and placed in the open field (45 × 45 × 20 cm^3^ [length × width × height]) (5 min test) 30 d after the first injection (i.e., paralleling the design from the contextual fear memory experiments).

#### Immunohistochemistry

Mice were perfused either 30 d after RV microinjection, 4 d following HSV microinjection, or 90 min after the memory retrieval event for c-Fos analysis. In all cases, mice were anesthetized with chloral hydrate and perfused intracardially with cold 0.1 M PBS followed by cold 4% PFA. Brains were removed, post-fixed in 4% PFA for 24 h and stored in a 30% sucrose solution until processed further. Brains were sectioned coronally on a cryostat (Leica CM1850), and 50-µm sections were obtained from throughout the hippocampus.

For GFP+DAPI staining, free-floating sections were washed (0.1M PBS), maintained in blocking solution (5% normal goat serum and 0.3% Triton X-100) for 2 h, then incubated with chicken anti-GFP (1:500; Aves Cat no. GFP-1020 RRID:AB_10000240) primary antibody for 72 h at 4 °C. Sections were washed (0.1M PBS) and incubated in Alexa 488 goat anti-chicken (1:500; Thermo Fisher Scientific Cat no. A-11039 RRID:AB_2534096) secondary antibody for 24 h at 4 °C. Sections were counterstained with DAPI (1:10000; Sigma, D9542) for 10 min, washed (0.1M PBS), then mounted on slides and coverslipped using Vectashield fluorescent mounting medium (Vector Laboratories).

For DCX+DAPI staining, sections were similarly maintained in blocking solution (4% normal donkey serum and 0.4% Triton X-100) for 2 h, then incubated with goat anti-DCX (1:500; Santa Cruz Cat no. sc-8066, RRID:AB_2088494) for 72 h at 4 °C. Sections were washed and incubated in Alexa 488 donkey anti-goat (1:500; Thermo Fisher Scientific Cat no. A-11055 RRID:AB_2534102) for 24 h at 4 °C.

For c-Fos+DAPI staining, sections were washed in 0.1 M PBS, maintained in blocking solution (5% normal goat serum and 0.3% Triton X-100) for 2 h, then incubated with rabbit anti-c-Fos (1:500; Synaptic Systems, 226 003 RRID:AB_2231974) for 72 h at 4 °C. Slices were washed (0.1 M PBS) and incubated in Alexa 633 goat anti-rabbit (1:500; Thermo Fisher Scientific A- 21071 RRID:AB_2535732) for 24 h at 4°C.

tdTomato and mCherry expression in Arc-TRAP mice and mice expressing AAV-RAM-hM3D_q_- mCherry was sufficiently strong such that amplification by immunohistochemistry was not required.

For BrdU+NeuN staining, free-floating sections were washed in 0.1 M PBS and then placed into 2N HCl heated at 45°C for 30 min. Sections were rinsed (0.1 M PBS) and then maintained in blocking solution (5% normal goat serum and 0.3% Triton X-100) for 2 h. Sections were then incubated with rat anti-BrdU (1:500; Bio Rad Cat No. MCA2060T RRID:AB 10015293) and NeuN (1:1000; Millipore Cat No. MAB377 RRID:AB_2298772) for 72 h at 4 °C. Slices were washed and incubated in Alexa 488 goat anti-rat (1:500; Thermo Fisher Scientific Cat No. A- 11006 RRID:AB_2534074) and Alexa 588 goat anti-mouse secondary antibodies (1:500; Thermo Fisher Scientific A-11031 RRID:AB_144696) for 24 h at 4 °C.

### Morphological analysis

To measure dendritic length, z-series at 0.5-µm intervals were acquired with a 20X lens with LSM710 confocal system (Zeiss). All cells were traced using Simple Neurite Tracer plugin for Fiji software. Dendritic complexity was quantified using the Sholl Analysis plugin for Fiji^85^.

To quantify spine density, we acquired GFP-labeled dendritic processes at the outer molecular layer using z-stacks with 0.5 µm intervals (objective 40X/1.3 NA; voxel size 0.07 × 0.07 × 1 µm; image size 1024 × 1024 pixels; 3X digital zoom). The number of spines were counted manually from the z-stack projections. Spine density was calculated by dividing the total number of spines by the length of the dendritic segment.

To quantify LMT density, high-resolution stacks (objective 40X/1.3 NA; voxel size 0.21 × 0.21 µm; image size 1024 × 1024 pixels) were acquired with LSM710 confocal system (Zeiss). Axonal length and number of LMTs (large, irregularly-shared swellings on mossy fibers in area CA3) were counted manually from the z-stack projections. LMT density was calculated by dividing the total number of LMTs by the length of the axonal segment. LMT size was calculated by tracing around the perimeter of LMTs and using Fiji software to calculate the area.

### Cell counts

BrdU and DCX were quantified to determine whether our gene deletion and optogenetic experiments affected overall rates of neurogenesis. Cells were counted using a 40X objective on an epifluorescent microscope. Cells were counted through the anterior–posterior extent of the DG using a 1/6 sampling fraction.

tdTomato, c-Fos, and DAPI sections were imaged on a laser confocal microscope (Zeiss LSM 710). All images were taken at 20X magnification. Frame size (512 × 515 pixels), image size (423 mm × 423 mm), and pixel size (0.42 mm) were kept consistent across groups and neural regions examined. Four images of dorsal DG (upper blade), CA3, and CA1 (bregma 1.46-2.18 mm) were analyzed per mouse. For each image, an optical z stack series and five to eight z- planes were counted for each neural region of interest within each mouse. Within each stack, the total number of DAPI^+^, tdT^+^, and c-Fos^+^ neurons were counted manually. The probability of neurons ‘tagged’ during memory encoding was calculated as *P*(tdT^+^|DAPI^+^). The probability of neurons activated during memory recall calculated as *P*(c-Fos^+^|DAPI^+^). Probability of engram reinstatement (i.e., tagged cells that were reactivated during memory recall test) was calculated as *P*(c-Fos^+^|tdT^+^). Chance levels of tdT reactivation was calculated as [(tdT^+^/DAPI^+^) × (c- Fos^+^/DAPI^+^)]. To determine whether tdT^+^ neurons were reactivated at levels greater than chance, we compared engram reinstatement to expected by chance. All counts and analysis were done using Fiji. mCherry^+^c-Fos^+^DAPI counts were performed identically to the tdT^+^c- Fos^+^DAPI counts.

### Statistical analysis

Morphological, cell count, and behavioral data were analyzed using either independent samples t-tests or mixed ANOVAs. Post-hoc LSDs were performed when appropriate. In one case, we used planned t-tests, based on strong a priori predictions (Figure 10). Statistical significance was set at *P* < 0.05.

## ACKNOWLEDGMENTS

This work was supported by Canadian Institutes of Health Research (CIHR) grants to PWF (FDN143227) and SAJ (MOP74650), and National Eye Institute and an RPB-Career Development Award to XD. AG was supported by fellowships from NSERC and SickKids.

## AUTHOR CONTRIBUTIONS

AG and PWF conceived the project. AG, JRE, SAJ, and PWF designed the experiments. AG and JD performed stereotaxic surgeries. EC and JS prepared retroviruses. AG, JdlP, SYK and JRE conducted the behavioural experiments. AG and JD conducted immunohistochemistry. AG, JdlP, MdS, and MS quantified neuron morphology. AG and PP conducted cell counts. AG conducted the statistical analysis. XD and JRS generated the conditional Cdh9 mice. PWF, SAJ, and AG wrote and edited the manuscript.

---

**Extended Data Figure 1.**
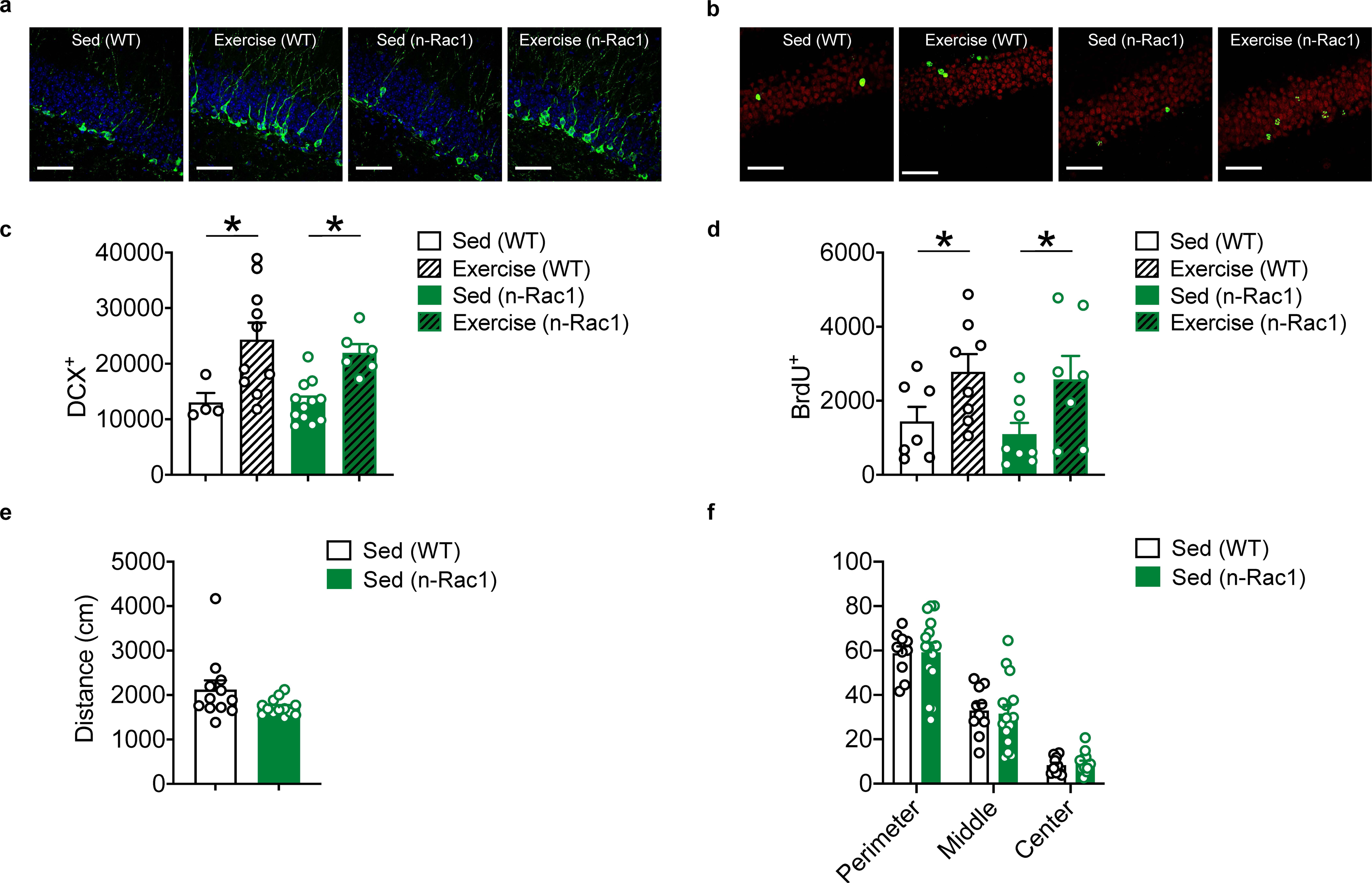
Adult neurogenesis and anxiety-like behaviors in n-Rac1 mice. Separate cohorts of WT and n-Rac1 mice were treated with the thymidine analog, BrdU, and TAM, and then given home cage access to a running wheel for 30 d (‘Exercise’ group) or housed conventionally (‘Sed’ group). Thirty d following TAM treatment, the number of BrdU^+^ cells and cells expressing the immature neuronal marker, doublecortin (DCX) were assessed in DG. Representative images illustrating (**a**) DCX^+^ and (**b**) BrdU^+^ cells. (**c**) Exercise increased numbers of DCX^+^ cells in both WT and n-Rac1 mice (ANOVA, no main effect of genotype: *F*_1,28_ = 0.25, *P* = 0.62; main effect of exercise: *F*_1,28_ = 17.54, *P* < 0.001; no genotype × exercise interaction: *F*_1,28_ = 0.23, *P* = 0.68). (**d**) Exercise increased numbers of BrdU^+^ cells in both WT and n-Rac1 mice (ANOVA, no main effect of genotype: *F*_1,26_ = 0.36, *P* = 0.56; main effect of exercise: *F*_1,26_ = 9.36, *P* < 0.005; no genotype × exercise interaction: *F*_1,26_ = 0.02, *P* = 0.88). (**e**) Distance travelled in open field did not differ (*t*_23_ = 1.91, *P* = 0.07). (**f**) Percent time spent in different open field zones did not differ between WT and n-Rac1 mice (ANOVA, no main effect of genotype: *F*_1,23_ = 0.01, *P* = 0.099, main effect of zone, *F*_2,46_ = 423.3, *P* < 0.001, no genotype × exercise interaction, *F*_2,46_ = 0.26, *P* = 0.77). Scale bars = 50 µm.

**Extended Data Figure 2.**
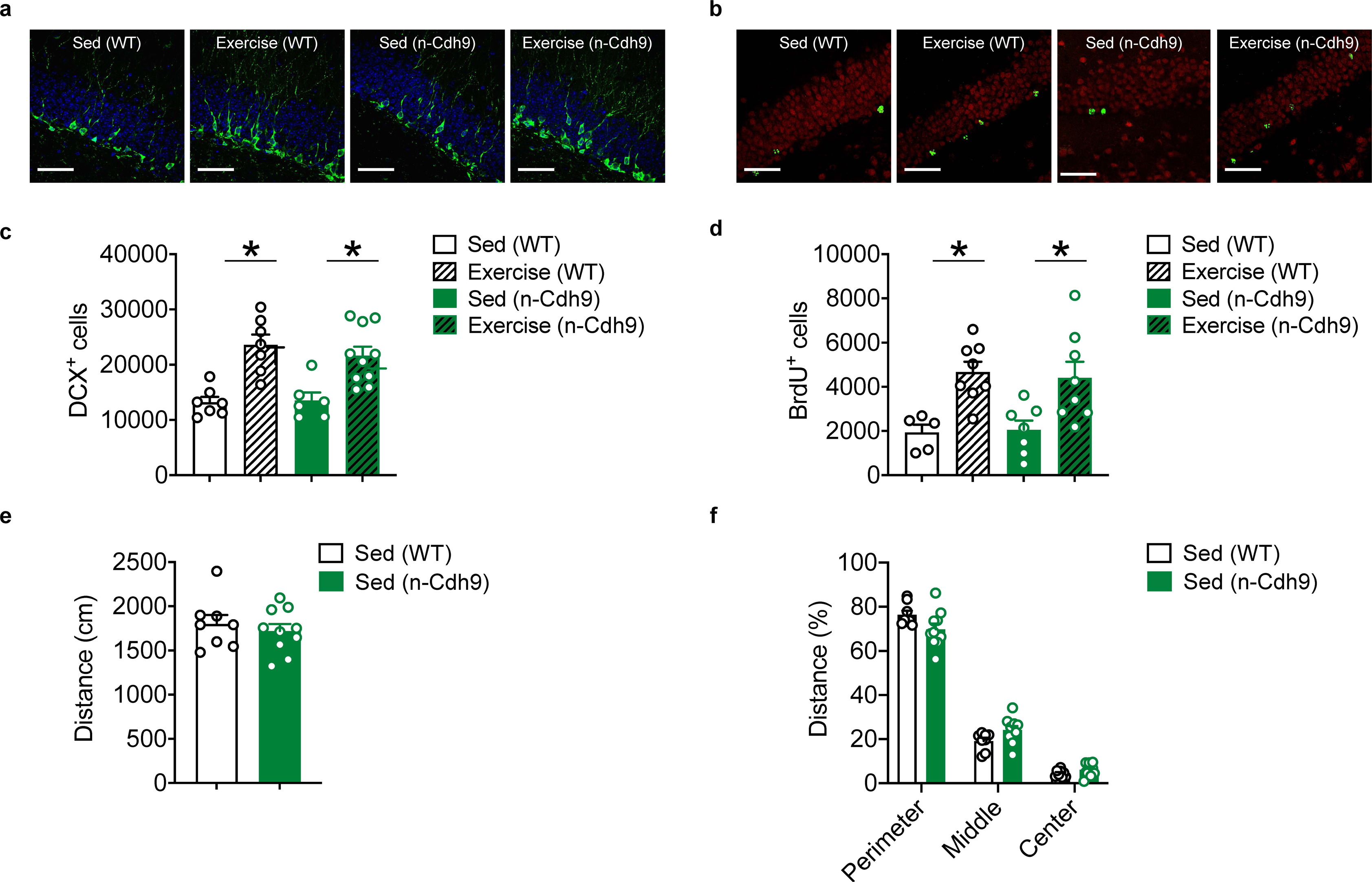
Adult neurogenesis and anxiety-like behaviors in n-Cdh9 mice. Separate cohorts of WT and n-Cdh9 mice were treated with the thymidine analog, BrdU, and TAM, and then given home cage access to a running wheel for 30 d (‘Exercise’ group) or housed conventionally (‘Sed’ group). Thirty d following TAM treatment, the number of BrdU^+^ and DCX^+^ cells were assessed in DG. Representative images illustrating (**a**) DCX^+^ and (**b**) BrdU^+^ cells. (**c**) Exercise increased numbers of DCX^+^ cells in both WT and n-Cdh9 mice (ANOVA, no main effect of genotype: *F*_1,28_ = 0.25, *P* = 0.62; main effect of exercise: *F*_1,28_ = 17.54, *P* < 0.001; no genotype × exercise interaction: *F*_1,28_ = 0.23, *P* = 0.68). (**d**) Exercise increased numbers of BrdU^+^ cells in both WT and n-Cdh9 mice (ANOVA, no main effect of genotype: *F*_1,24_ = 0.26, *P* = 0.61; main effect of exercise: *F*_1,24_ = 11.90, *P* < 0.005; no genotype × exercise interaction: *F*_1,24_ = 0.74, *P* = 0.40). (**e**) Distance travelled in open field did not differ (*t*_16_ = 0.62, *P* = 0.18). (**f**) Percent time spent in different open field zones did not differ between WT and n-Cdh9 mice (ANOVA, no main effect of genotype: *F*_1,16_ = 1.98, *P* = 0.18, main effect of zone, *F*_2,32_ = 539.2, *P* < 0.0001; genotype × exercise interaction, *F*_2,32_ = 3.82, *P* < 0.05). Scale bars = 50 µm.

**Extended Data Figure 3.**
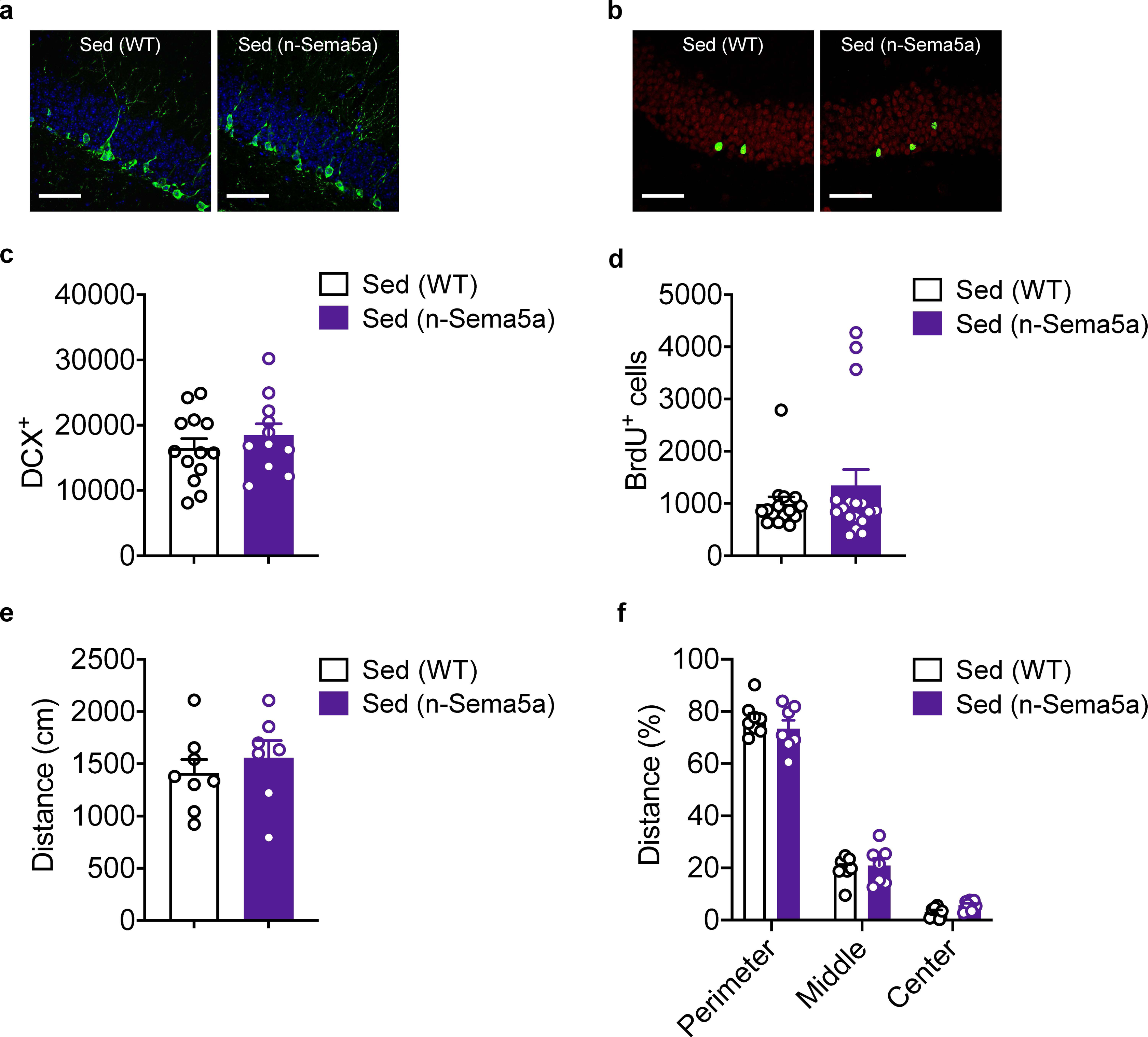
Adult neurogenesis and anxiety-like behaviors in n-Sema5A mice. Separate cohorts of WT and n-Sema5A mice were treated with the thymidine analog, BrdU, and TAM. Thirty d following TAM treatment, the number of BrdU^+^ and DCX^+^ cells were assessed in DG. Representative images illustrating (**a**) DCX^+^ and (**b**) BrdU^+^ cells. (**c**) Similar numbers of DCX^+^ cells in WT and n-Sema5A mice (*t*_22_ = 0.89, *P* = 0.39). (**d**) Similar numbers of BrdU^+^ cells in WT and n-Sema5A mice (*t*_30_ = 1.00, *P* = 0.32). (**e**) Distance travelled in open field did not differ (*t*_13_ = 0.71, *P* = 0.49). (**f**) Percent time spent in different open field zones did not differ between WT and n-Cdh9 mice (ANOVA, no main effect of genotype: *F*_1,13_ = 3.17, *P* = 0.10, main effect of zone, *F*_2,26_ = 433.5, *P* < 0.0001, no genotype × exercise interaction, *F*_2,26_ = 0.85, *P* = 0.44). Scale bars = 50 µm.

**Extended Data Figure 4.**
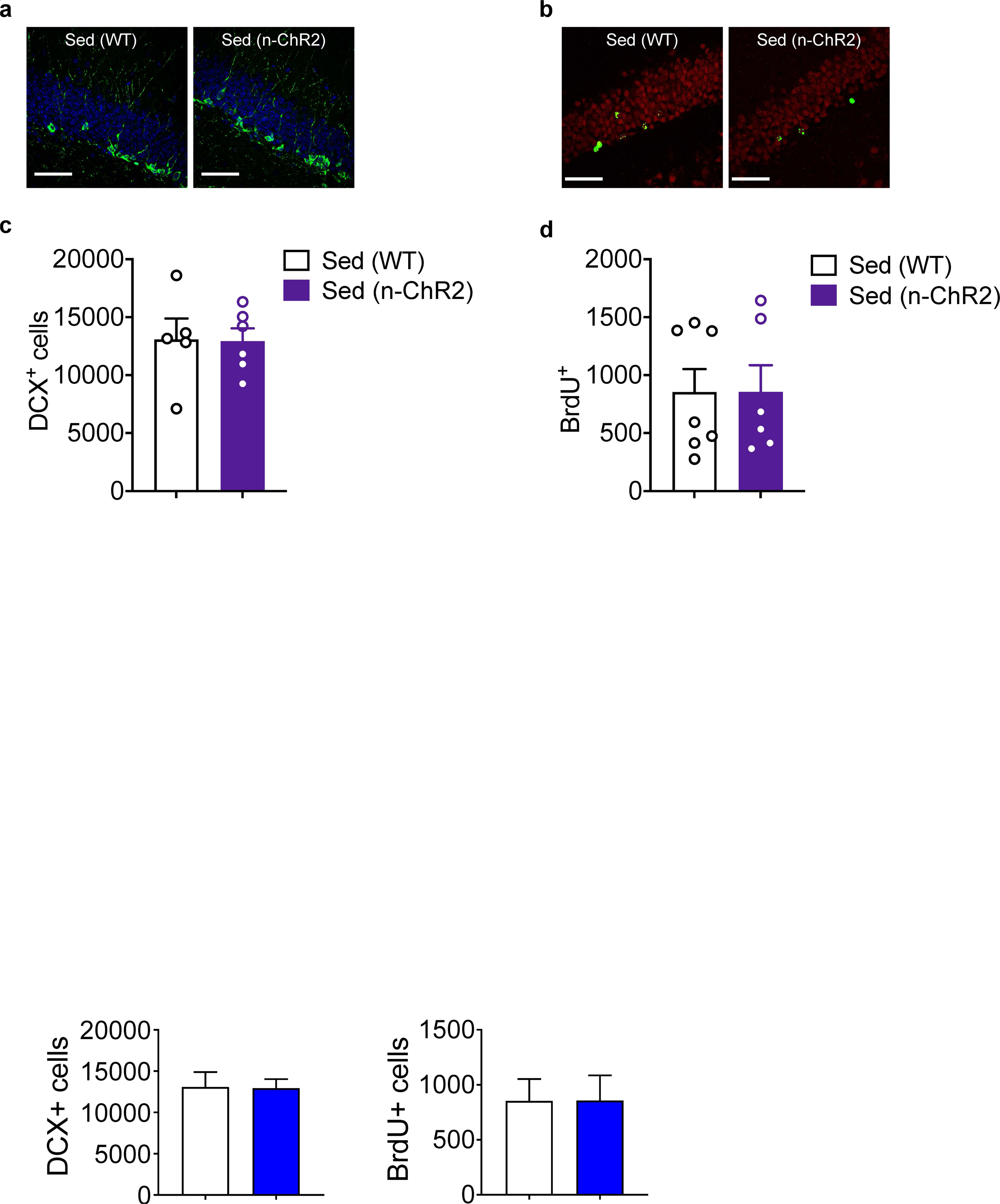
Adult neurogenesis in n-ChR2 mice. Separate cohorts of WT and n- Sema5A mice were treated with the thymidine analog, BrdU, and TAM. Thirty d following TAM treatment, the number of BrdU^+^ and DCX^+^ cells were assessed in DG. Representative images illustrating (**a**) DCX^+^ and (**b**) BrdU^+^ cells. (**c**) Similar numbers of DCX^+^ cells in WT and n-ChR2 mice (*t*_9_ = 0.06, *P* = 0.95). (**d**) Similar numbers of BrdU^+^ cells in WT and n-ChR2 mice (*t*_11_ = 0.01, *P* = 0.99). Scale bars = 50 µm.

**Extended Data Figure 5.**
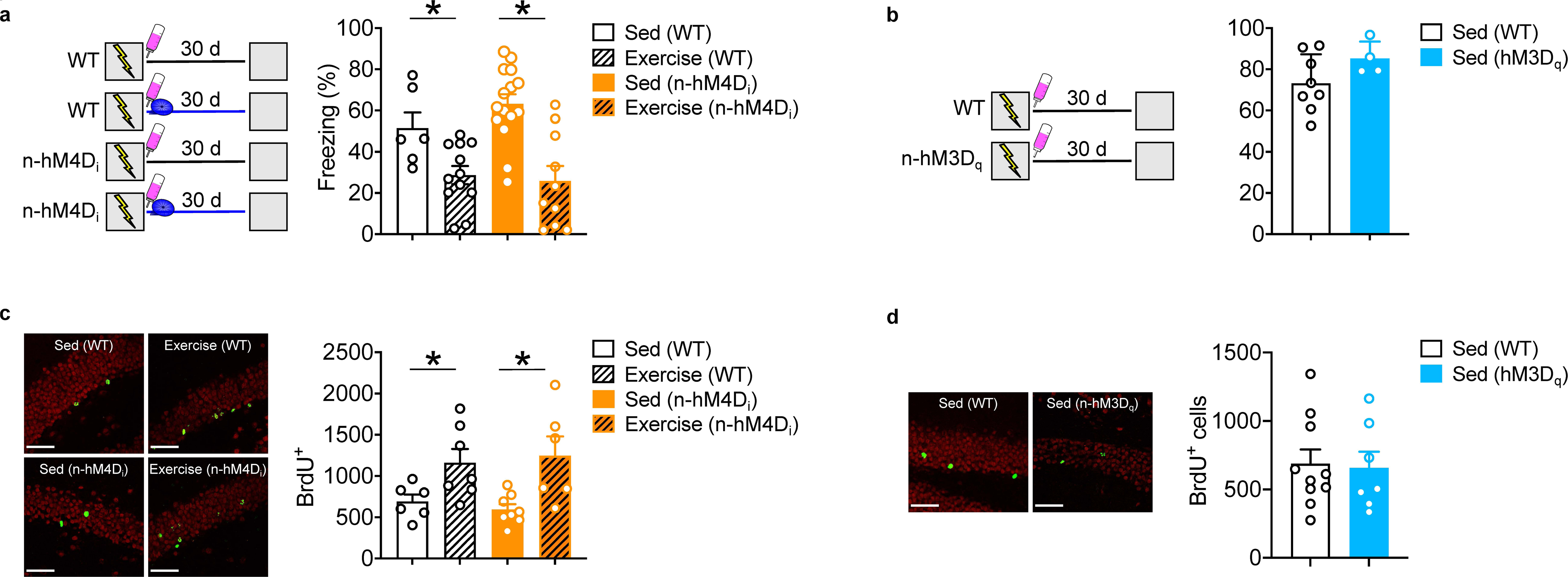
Behavioral controls and analysis of global neurogenesis in n-hM4D_i_ and n-hM3D_q_ mice. (**a**) (left) Experimental design. n-hM4D_i_ mice were trained in contextual fear conditioning. Mice either had access to an exercise wheel (‘Exercise’ groups) or were housed conventionally (‘Sed’ groups), and then were tested 30 d later. In this experiment, mice were not treated with TAM. Mice received CNO (via drinking water) throughout the retention period. (right) Exercise reduced freezing in both WT and n-hM4D_i_ mice (ANOVA, no main genotype effect: *F*_1,39_ = 0.54, *P* = 0.47; main exercise effect: *F*_1,39_ = 24.99, *P* < 0.001; no genotype × exercise interaction: *F*_1,39_ = 1.46, *P* = 0.24). (**b**) (left) Experimental design. n-hM3D_q_ mice were trained in contextual fear conditioning, and tested 30 d later. In this experiment mice were not treated with TAM. Mice received CNO (via drinking water) throughout the retention period. (right) Freezing was similar in n-hM3D_q_ compared to WT mice (*t*_10_ = 1.56, *P* = 0.15). (**c**) Experimental design. n-hM4D_i_ mice were treated with TAM (to induce expression of hM4D_i_ in NPCs and their progeny) and BrdU (to label dividing cells) and then had access to an exercise wheel (‘Exercise’ groups) or were housed conventionally (‘Sed’ groups) for 30 d. Mice received CNO (via drinking water) throughout the 30 d delay. (right) Exercise increased neurogenesis in both WT and n-hM4D_i_ mice (ANOVA, no main genotype effect: *F*_1,23_ = 0.002, *P* = 0.97; main exercise effect: *F*_1,22_ = 14.85, *P* < 0.001; no genotype × exercise interaction: *F*_1,22_ = 0.40, *P* = 0.53). (**d**) (left) Experimental design. n-hM3D_q_ mice were treated with TAM (to induce expression of hM4D_i_ in NPCs and their progeny) and BrdU (to label dividing cells). Mice received CNO (via drinking water) for 30 d, and the numbers of BrdU^+^ cells were assessed. (Right) There were similar numbers of BrdU+ cells in WT vs. n-hM3Dq mice (*t*_15_ = 0.19, *P* < 0.85). Scale bars = 50 µm.

